# DAMEfinder: A method to detect differential allele-specific methylation

**DOI:** 10.1101/800383

**Authors:** Stephany Orjuela, Dania Machlab, Mirco Menigatti, Giancarlo Marra, Mark D. Robinson

## Abstract

DNA methylation is a highly studied epigenetic signature that is associated with regulation of gene expression, whereby genes with high levels of promoter methylation are generally repressed. Genomic imprinting occurs when one of the parental alleles is methylated, i.e, when there is inherited allele-specific methylation (ASM). A special case of imprinting occurs during X chromosome inactivation in females, where one of the two X chromosomes is silenced, in order to achieve dosage compensation between the sexes. Another more widespread form of ASM is sequence dependent (SD-ASM), where ASM is linked to a nearby heterozygous single nucleotide polymorphism (SNP).

We developed a method to screen for genomic regions that exhibit loss or gain of ASM in samples from two conditions (treatments, diseases, etc.). The method relies on the availability of bisulfite sequencing data from multiple samples of the two conditions. We leverage other established computational methods to screen for these regions within a new R package called DAMEfinder. It calculates an ASM score for all CpG sites or pairs in the genome of each sample, and then quantifies the change in ASM between conditions. It then clusters nearby CpG sites with consistent change into regions.

In the absence of SNP information, our method relies only on reads to quantify ASM. This novel ASM score compares favourably to current methods that also screen for ASM. Not only does it easily discern between imprinted and non-imprinted regions, but also females from males based on X chromosome inactivation. We also applied DAMEfinder to a colorectal cancer dataset and observed that colorectal cancer subtypes are distinguishable according to their ASM signature. We also re-discover known cases of loss of imprinting.

We have designed DAMEfinder to detect regions of differential ASM (DAMEs), which is a more refined definition of differential methylation, and can therefore help in breaking down the complexity of DNA methylation and its influence in development and disease.

## Background

Epigenetic modifications refer to mitotically-heritable, chemical variations in DNA and chromatin in the absence of changes in the DNA nucleotide sequence itself [1, 2]. Although there are a large number of such documented phenomena, DNA methylation (i.e., methyl groups added to cytosines in mammalian DNA, mostly in CpGs dinucleotides) stands out because the mechanism of heritability, via maintenance methyltransferases, is well-determined [3–5]. In addition, due to well-known effects of chemical reactions, such as sodium bisulfite conversion of cytosines to uracils [6], and biochemical reactions like TET-pyridine borane conversion of 5-methylcytosine to dihydrouracil [7], the interrogation of DNA methylation level across the genome can be sampled and quantified at each cytosine.

DNA methylation plays a role in several biological phenomena. It is believed to be associated with gene expression, with the canonical relationship suggesting that transcriptional units with high levels of promoter methylation are repressed or silenced, although not all genes with unmethylated promoters are switched on, since other epigenetic mechanisms of silencing may come into play [8].

Genomic imprinting, where genes are expressed in a parent-of-origin manner [9], is also regulated by DNA methylation. Imprinting occurs via allele-specific methylation (ASM), in which only the paternal or the maternal allele is methylated in all or most of the tissues of an individual [9]. This methylation asymmetry is conferred during gametogenesis in the parental germlines, or during early embryogenesis after fertilization, and will remain during the lifetime of the offspring [10]. A recent survey [11] reported 228 genes linked to imprinted control, and from those, 115 linked to imprinted regulation in human placenta. These genes are known for their roles in embryonic and fetal development, placental formation, cell growth and differentiation, metabolism and circadian clock regulation [11]. In fact, loss of imprinting and abnormal expression of imprinted genes are implicated in severe congenital diseases, like the neurodevelopment disorders Angelman and Prader-Willi syndromes. The first is caused by the lack of maternal *UBE3A* gene expression, the second by loss of paternal expression of several contiguous genes on chromosome 15q11-q13 [12]. Furthermore, disruption of imprinting in somatic cells has been implicated in the pathogenesis of different cancers, like loss of imprinting within the *H19/IGF2* imprinting control region in colorectal cancer [13], and gain of imprinting at 11p15 in hepatocellular carcinoma [14].

A special and well characterized case of imprinting occurs during X chromosome inactivation (XCI), where one of the two X chromosomes is randomly silenced via DNA methylation and other epigenetic mechanisms, early in development in each cell of a female, in order to achieve dosage compensation between the sexes [15].

Beside imprinting and XCI, the rest of the genome is thought to be symmetrically methylated across both alleles. However, sequence-dependent ASM (SD-ASM) has been frequently reported in the last 10 years and appears to be widespread in the human genome [16–21]. In this case, the DNA methylation asymmetry between the parental alleles appears to be causally related to the presence of a single nucleotide polymorphism (SNP). As for imprinted ASM and XCI, SD-ASM can be associated with silencing of one of the two parental gene copies, likely mediated by cis-acting, allele-specific changes in affinity of DNA-binding proteins [21]. Thus, SD-ASM would explain why a large number of genes are differently expressed among individuals in a given cell type. SD-ASM appears to be also tissue-specific [22, 23], thus it is commonly believed that the interaction between genetic variants (i.e., SNPs) and epigenetic mechanisms (i.e., effects of DNA methylation asymmetry on gene expression) modulates the susceptibility of the general population to frequent, multi-factorial diseases affecting specific organs, such as ASM in the *PEAR1* intron 1, which is linked to platelet reactivity and cardiovascular disease [24]; or ASM in *FKBP5* enhancers, which poses an increased risk to stress-related psychiatric disorders in individuals who suffered an abuse during childhood [25]. Although the modulation of the susceptibility to a complex disease by a SD-ASM is generally weak and influenced by environmental factors, it is worth noting that 5-10% of all SNPs might be associated with SD-ASMs in the genome of a given tissue of a given individual [19, 20, 26].

Although there are several technologies to study DNA methylation, such as microarrays that genotype bisulfite-converted DNA, or lower resolution capture technologies such as methyl-binding domain (MBD) sequencing [27], or methylated DNA immunoprecipitation (MeDIP) sequencing [28], bisulfite sequencing (BS-seq) remains distinct for the ability to read out DNA methylation of a single allele at base-resolution. Importantly, BS-seq can be conducted both in an unbiased genome-wide fashion, or in combination with technologies that focus the sequencing to particular regions, either by making use of hybridization or enzyme digestions [29].

Recent studies have obtained ASM readouts from mapped bisulfite reads, by assigning them to the alleles of each known heterozygous SNP. Methylation levels are then determined for all allele-linked cytosines in the reads (see [20, 30, 31] for recent examples). The ASM calculated in this way is interpreted as SD-ASM, and it does not include imprinted ASM nor XCI, since they are not necessarily sequence dependent. Calculating ASM in this fashion is limited by the availability of SNP information from either DNA-seq or SNP-array data, or directly from the BS-seq reads [32]. However performing different types of high-throughput experiments is economically restrictive and time consuming, and deriving SNPs from BS-seq reads can be problematic due to bisulfite conversion of DNA (i.e., distinguishing a C/T SNP from a C/T conversion of a methylated cytosine) and imbalanced strand coverage (i.e., when the Watson and Crick strands are not equally or highly covered) [32].

Considering these limitations in ASM detection, a couple of studies have sought to make sole use of BS-seq reads to screen for the full spectrum of ASM. The tools **allelicmeth** and **amrfinder** (from the same authors) [33] are the only available executable methods that detect ASM without SNP information. Briefly, the **allelicmeth** method creates a contingency table with the counts of methylated and unmethylated reads covering a pair of CpG sites. A score is calculated via Fisher’s exact test that represents the probability that both CpG sites have an equal proportion of methylated-unmethylated reads. **amrfinder** also calculates ASM but at a regional level. It fits two statistical models, one assuming that both alleles are equally methylated, and the other assuming different methylation states for the two alleles. A region is considered to have ASM by comparing the likelihoods of the two models. A more recent algorithm termed *MethylMosaic* relies on the principle that bimodal methylation patterns, independent from the genotype, are a good indicator of ASM [34]; however, to our knowledge there is no publicly available implementation.

Based on the current state of ASM detection from BS-seq reads, we set out to develop a simple yet effective method to screen for genomic regions that exhibit loss or gain of ASM between samples from distinct conditions. The methods mentioned above detect ASM in individual samples, however they do not allow a flexible comparison between groups of samples, such as that performed in a typical differential methylation analysis [35, 36], where the goal is to find the effect of treatments or diseases on methylation, reflected as increase or decrease of methylation levels. Here, we are interested in performing such differential analysis but focusing on the effect of ASM, reflected as gain or loss of allele-specificity. For this task, we introduce DAMEfinder (Differential Allele-specific MEthylation finder), an **R** package [37] that consists of: i. a scoring function that reflects ASM for several samples; ii. integration with **limma** [38] and **bumphunter** [39] to detect differentially allele-specific methylated regions (DAMEs); and, iii. accurate estimation of false discovery rates (FDR). We demonstrated the ASM score and DAMEfinder on two real data sets, one based on targeted-enrichment BS-seq, comparing normal colonic mucosa to cancerous colorectal lesions, and another on whole genome BS-seq (WGBS), comparing blood monocytes from healthy females and males.

## Results

### The overall DAMEfinder workflow

Figure 1 gives an overview of the pipeline. We make considerable use of existing tools and keep inputs/outputs in standard formats. In order to make use of the package, the user must independently use **bismark** to map paired-end BS-seq reads against a reference genome (Figure 1A). Once this is done, the user has the option to detect ASM for each sample in two ways: (1) Using the output from **methtuple** [40], which computes read counts of *pairs* of nearby CpG sites. From these counts, we compute an ASM score; and/or (2) using an additional VCF file containing heterozygous SNPs. For each SNP we call methylation from the reads containing that SNP, and calculate an ASM score for each CpG site (Figure 1B and details below). From the set of scores, we leverage routines from the **bumphunter** and **limma** packages to calculate a statistic and detect regions showing persistent change in ASM. We call these regions DAMEs (Figure 1C). We estimate and control a regional FDR through permutations or by implementation of the Simes method [41].

**Figure 1.**
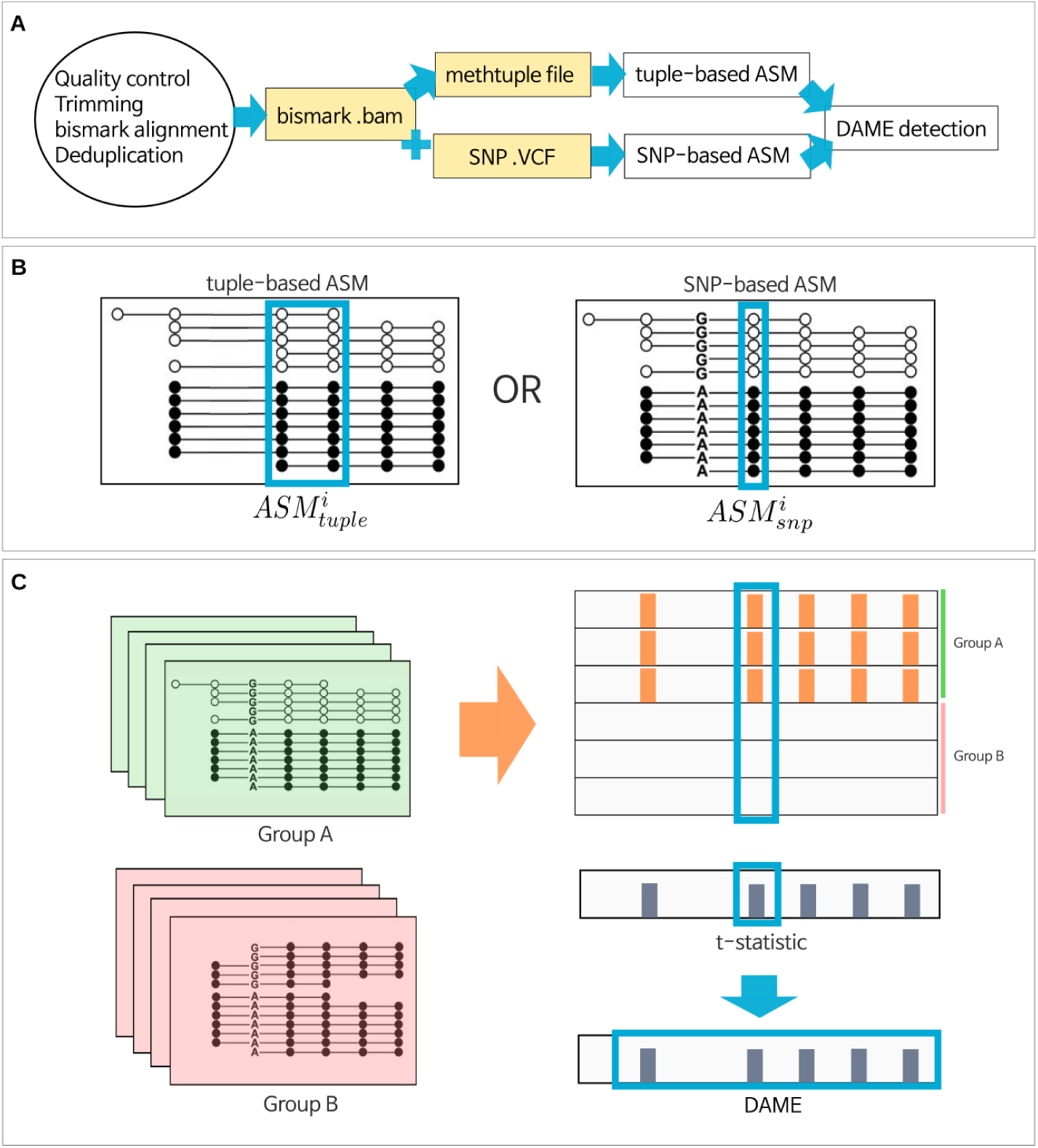
The DAMEfinder pipeline. **A.** Files necessary to run DAMEfinder are reported in yellow rectangles. White rectangles show the main R outputs from DAMEfinder. Steps to be run before DAMEfinder are in the circle, i.e., fastq files undergo quality control and read alignment with **bismark** [42]. The resulting bam file is used to calculate an ASM score, which can be done in two ways: **B.** (i) the tuple-based strategy that takes as input a beforehand created **methtuple** [40] file. The score is calculated based on the read counts of pairs of CpG sites. (ii) the SNP-based strategy, which takes as input both the bam file and a VCF file with heterozygous SNPs. Here the score is calculated for each CpG site in the reads containing a SNP. **C.** We determine differential ASM by calculating a statistic based on either the tuple ASM or the SNP-ASM (using **limma** [38]), which reflects the difference between two conditions (Group A vs. Group B) for each genomic position (tuple or site). DAMEs are defined based on this statistic, as regions of contiguous positions with a consistent change in ASM.

### The ASM score

#### SNP-based ASM

The most straightforward way of detecting ASM from mapped reads, is by assigning them to either of the alleles at each known heterozygous SNP. Methylation status is then determined for each allele-linked cytosines in the reads. We have used this strategy to calculate a SNP-based ASM score 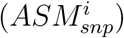, and consider it to be the genuine form of ASM, since it is derived from an extra layer of information, i.e. the genotype of an individual.

We extract the reads overlapping every heterozygous SNP in a VCF file with the **GenomicAlignments** R package [43], and for each read determine the methylation status of the CpG sites. Sites that are not in reads containing a SNP are not considered. We calculate 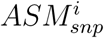 for each CpG site *i* contained in the reads of a SNP as:

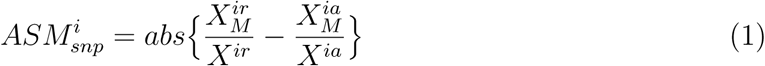

where 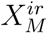 and 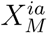 correspond to the number of methylated reads from the reference *r* allele, and the alternative *a* allele. In practice, it makes no difference which allele is the reference or the alternative. *X*^*ir*^ and *X*^*ia*^ correspond to the total number of reads covering the reference and the alternative allele (see schematic in Figure 1B). The score ranges from 0 to 1, where a score of 1 represents the scenario where one allele is completely methylated, and the other allele is fully unmethylated; a value of 0 means an equal proportion of methylated sites in both alleles.

#### Tuple-based ASM

Instead of restricting ASM detection to allele-linked reads, we can make use of an entire set of CpG sites to detect ASM. For this task, we designed a score under the assumption that pairs of CpG sites in the same DNA molecule (read) are correlated [44, 45], and that in a biallelic organism, intermediate levels of methylation could represent allele-specificity, i.e., the proportion of methylated reads in a pair of CpG sites or tuple is close to 0.5. We calculate this score as a weighted log-odds ratio:

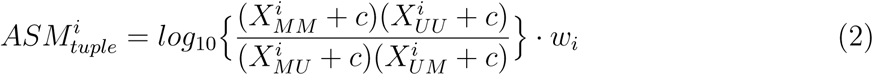

where 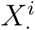 corresponds to the number of reads covering a unique pair of CpG sites *i*, generated by running the **methtuple** tool. CpG sites in a pair can be methylated *MM*, unmethylated *UU*, or mixed (*UM* or *MU*). A constant *c* is added to every *X*^*i*^ to avoid dividing by 0. The log-odds ratio is multiplied by a weight, *w*_*i*_, which is set such that the ratio of *MM* :*UU* can depart somewhat from a 50:50 relation, while *MM* or *UU* tuples, which represent absence of allele-specificity, are attenuated to 0. This is calculated as:

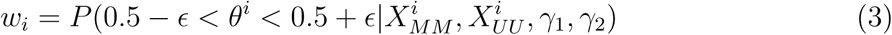

where *ϵ* represents the degree of allowed departure from a 50:50 ratio, and *θ*^*i*^:

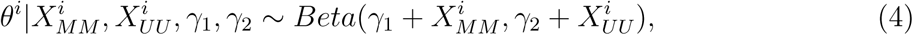

represents the moderated proportion of *MM* to *MM* +*UU* reads. It is based on a beta model, where *γ*_1_ and *γ*_2_ are hyperparameters set to penalize fully methylated or fully unmethylated tuples, i.e., when the *MM* : *UU* balance goes farther from a 50:50 relation. Similar to 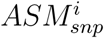, higher values of 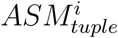 (can be higher than 1), indicate putative presence of allele-specificity.

### ASM score validation

In order to test the *ASM*_*tuple*_ score, we used the *ASM*_*snp*_ score as an indicator of true ASM, and calculated the *ASM*_*tuple*_ score, the **allelicmeth** and **amrfinder** scores, and a score representing absolute deviation from 50% methylation (methdeviation; see Methods), in a single normal tissue sample from the colorectal cancer (CRC) dataset (see Methods).

Figure 2 shows the true positive rate (TPR) and false positive rate (FPR) achieved by the 4 evaluated scores at 3 different coverage thresholds (left to right), and 2 *ASM*_*snp*_ cutoffs (top to bottom). *ASM*_*tuple*_ was consistently more sensitive and specific than the other three scores, especially as coverage was increased. Intermediate methylation values yielded comparable results, however the *ASM*_*tuple*_ was able to detect more cases of “real” ASM in all combinations. **allelicmeth** increasingly failed as coverage and *ASM*_*snp*_ value increases. **amrfinder** performed better than **allelicmeth** at higher true values.

**Figure 2.**
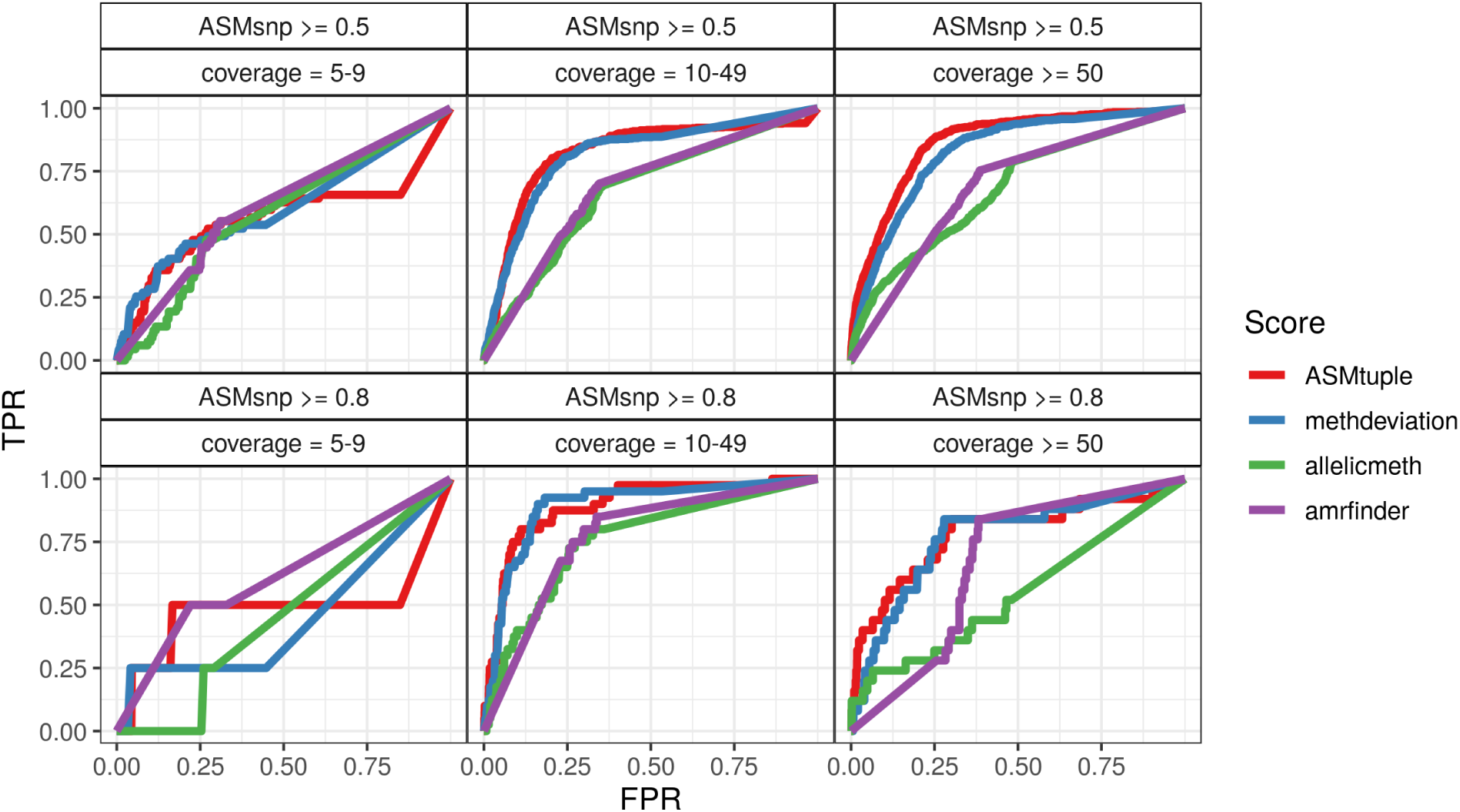
Comparison of the *ASM*_*tuple*_ score to **allelicmeth, amrfinder** and methylation deviation, by considering *ASM*_*snp*_ as true ASM. We calculated *ASM*_*tuple*_ scores (red), deviations from 50% methylation (blue), **allelicmeth** scores (green), **amrfinder** scores (purple) in a sample of normal colorectal mucosa included in the CRC dataset. The scores were compared to each other by plotting the FPR against the TPR achieved. The plots are drawn for different intervals of read coverage (5-9, 10-49, ≥ 50), and different levels of the *ASM*_*snp*_ score (≥ 0.5, ≥ 0.8), which is considered the “true” ASM. Overall AUCs (area under the curve) for the top three panels: *ASM*_*tuple*_ = 0.83, deviations from 50% = 0.81, **allelicmeth** = 0.66, **amrfinder** = 0.68. Overall AUCs for the lower three panels: *ASM*_*tuple*_ = 0.82, deviations from 50% = 0.81, **allelicmeth** = 0.64, **amrfinder** = 0.72

As an additional validation of the *ASM*_*tuple*_ score, we used the blood dataset (see Methods) to compare healthy male and female samples. In principle, females should exhibit allele-specificity in the X chromosome due to XCI and thus higher *ASM*_*tuple*_ values. Figure 3 shows the distribution of *ASM*_*tuple*_ values across all samples in the dataset, in chromosome 3 and chromosome X. From a whole genome perspective (Figure 3A), there is little difference between males and females in X chromosome (mean of row-means females: 0.13, males: 0.098), and practically no difference in chromosome 3 (0.060, 0.074). However, by focusing on CpG tuples located in promoter regions (1 kb upstream the transcription start site - TSS), we observed ASM values increased only in chromosome X of females (Figure 3B; 0.30, 0.088).

**Figure 3.**
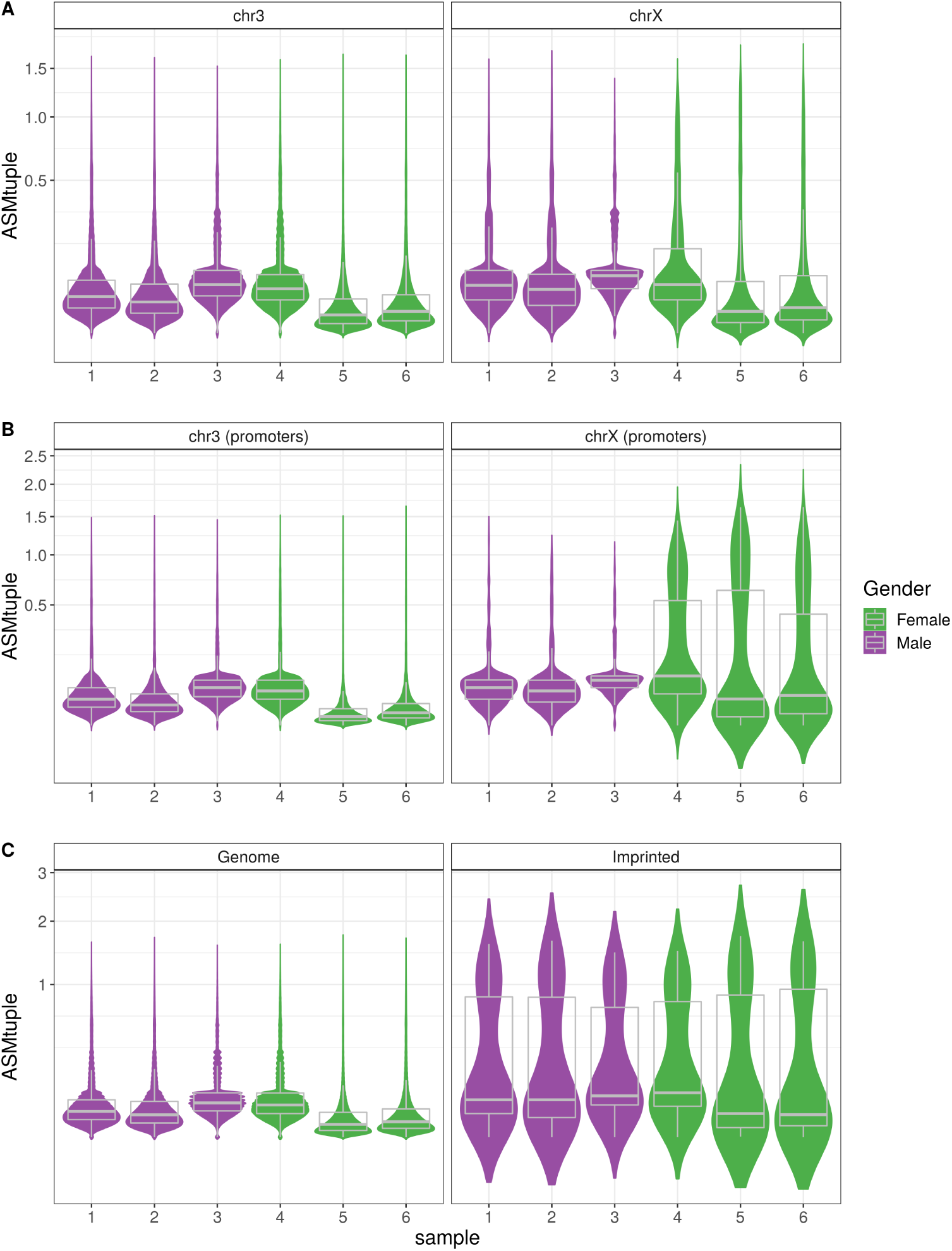
*ASM*_*tuple*_ distribution in the genome. We used XCI as a proof of concept for allele-specificity in females. Data from the blood dataset comprising 3 females and 3 males was used for this analysis. **A**. When considering all CpG tuples in the genome, the *ASM*_*tuple*_ distribution (y-axis) in chromosome 3 and chromosome X is similar in both genders. **B**. When considering CpG tuples located in promoter regions (i.e., 1 kb upstream of the TSS), the *ASM*_*tuple*_ score is higher in chromosome X of females. **C**. Promoter regions of 89 known imprinted regions (see Methods) also exhibit higher *ASM*_*tuple*_ compared to values in the rest of the genome. Y-axis in all plots is square-root transformed

In the same blood dataset, we also compared the *ASM*_*tuple*_ scores from the promoters of imprinted genes reported in [11] (see Methods), to the scores from rest of the genome (Figure 3C). As expected, ASM scores were higher in the tuples located within imprinted promoters, for both males and females.

### DAME detection

As depicted in Figure 1, after calculating *ASM*_*tuple*_ or *ASM*_*snp*_ in the DAMEfinder pipeline, we continue to detect regions of persistent change in ASM between one condition to another within a cohort of samples. Change can occur as loss of ASM, when a reference group exhibits allele-specificity across a region (high values of ASM), and the group of interest has this same region fully methylated, unmethylated, or with random methylation (low values of ASM). Change can also occur as gain of ASM, where the reference group does not have allele-specificity and the group of interest does. We call regions such as this DAMEs (Differentially Allele-specifically MEthylated regions).

To detect DAMEs, we first obtain a regression coefficient *β*_*ij*_ followed by a t-statistic using the R package **limma** [38] (see Methods), on the transformed 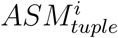 score, or on the 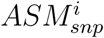 score, for each CpG position *i* (tuple or site), across *j* samples (see Methods for model).

We detect regions of contiguous CpG positions where *β*_*ij*_ persistently deviates in the same direction from zero; this is done in two ways:

#### Permuting bumphunted-regions

The **regionFinder** function from **bumphunter** is used to scan for regions (*R*) where CpG sites close in proximity have *β*_*ij*_ above a user-defined threshold *K*, which corresponds to a percentile of *β*_*ij*_. For each region detected, the function also calculates an area A= ∑_*iϵR*_ |*β*_*ij*_|. For the CRC data set, we used the default value *K* = 0.7, and distance between CpG positions up to 100 bp.

We assess significance of every region detected by assigning an empirical p-value. For every non-redundant, permutation of the coefficient of interest (chosen from a column in the design matrix *X*), **regionFinder** is applied again. All the areas generated by all permutations are pooled to generate a null distribution of areas [46]. We define the p-values for each *R* as the proportion of null areas greater than the observed *A*; p-values are adjusted using the Benjamini-Hochberg method [47] from the **stats** R package [37].

#### Cluster-wise correction

Optionally, we define regions that exhibit changes in ASM by first generating clusters of CpG sites with **clusterMaker**. For each cluster, we aggregate all the CpG position p-values generated by **limma** using the Simes method [41], which is applicable when test statistics exhibit positive dependence [48]. As implemented in [49], we calculate:

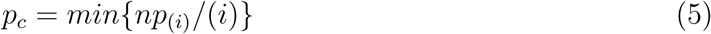

where *p*_(1)_, …, *p*_(*n*)_ are the ordered p-values of each CpG position *i* in a cluster *c* and *n* is the number of CpG positions in the cluster. *p*_*c*_ summarizes evidence against the null hypothesis that all CpG positions are not differential. We adjust *p*_*c*_ as above.

#### Evaluation of DAME detection

We compared the different strategies to control FDR in the DAME detection pipeline, by applying them to a semi-simulated dataset and plotting the TPR and FDR achieved at different adjusted p-value thresholds (0.01, 0.05, 0.1) (Figure 4). We designed a small set of simulated DAMEs to evaluate the FDR control of the above strategies. We took 6 samples of normal tissue from the CRC dataset and calculated *ASM*_*snp*_ scores in each of them. We assumed these scores to be the *ASM*_*snp*_ baseline in the simulation. Then, we divided the samples into two groups of three samples each, and for all the CpG sites covered by the 6 samples, we defined clusters of contiguous CpG sites. For each truly differential cluster, we added signal to a randomly determined subset of adjacent CpG sites (see Methods for more details).

**Figure 4.**
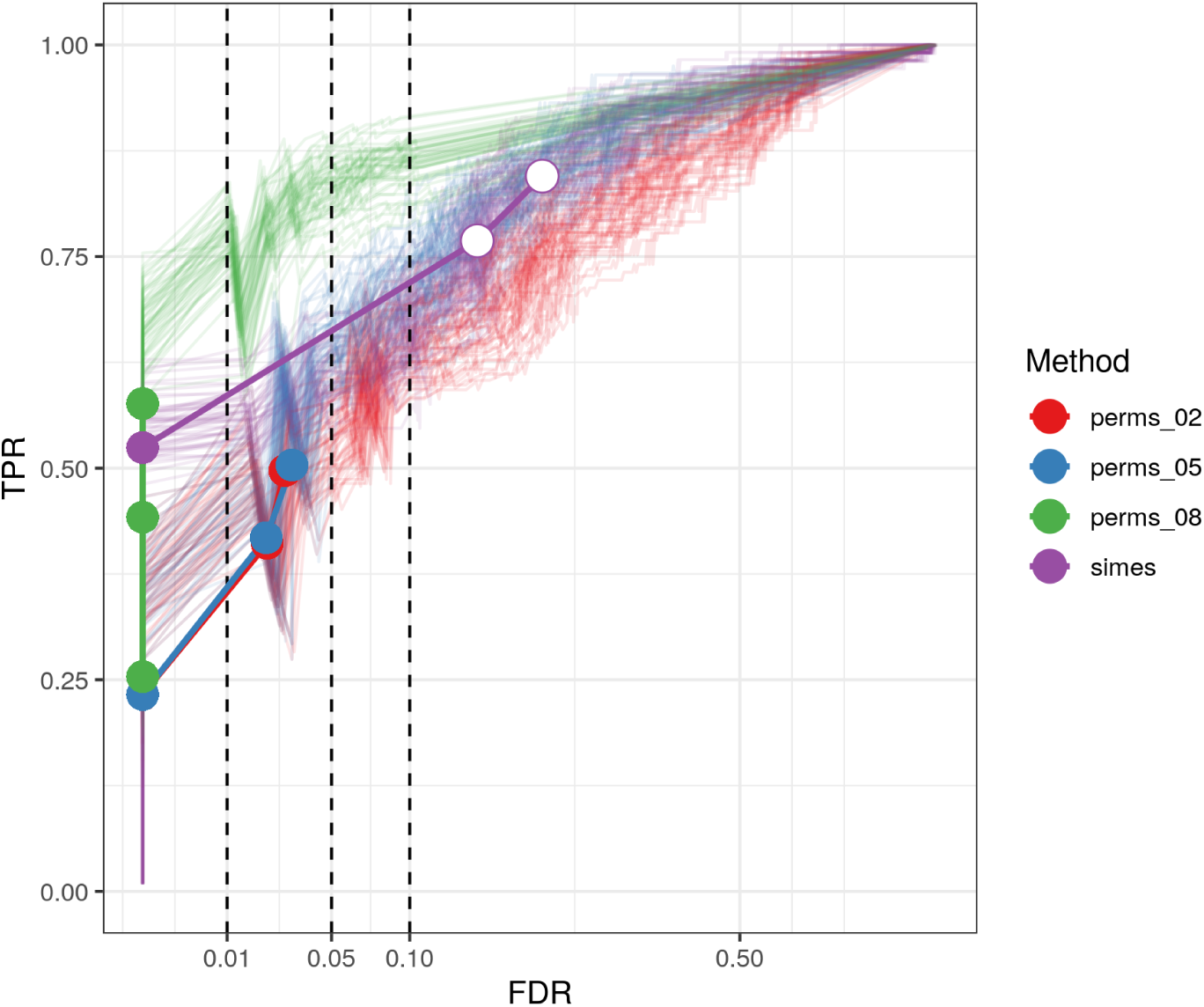
FDR control of p-value assignment strategies. We plot the FDR against the TPR achieved by the two alternatives for assigning p-values to a DAME: The first by generating permutations and setting a threshold *K* (see text) on the t-statistic (here 0.2,0.5,0.8), the second by using the Simes method. Lines are colored by strategy. Each strategy was run 50 times with the same simulation parameters. Colored circles indicate that the FDR achieved is smaller than the specified threshold (dashed lines at 0.01, 0.05 and 0.1), white circles indicate the opposite. x-axis is square-root transformed.

Overall, the empirical p-value controlled the FDR, whereas the Simes method tended to be less conservative but more sensitive (Figure 4 and Supplementary Figure 1, Additional File 1 for same plot tested with different parameters).

### Discovery of DAMEs in colorectal cancer dataset

We used a previously published dataset comprising 6 patients with diagnosed colorectal cancer, three with CIMP (CpG-Island Methylator Phenotype), and three without CIMP (see Methods); DNA from normal mucosa and cancer lesions was bisulfite-sequenced. We ran **DAMEfinder** on this dataset in both modes, therefore obtaining the *ASM*_*snp*_ and *ASM*_*tuple*_ scores. After filtering for coverage (more than 5 reads) and for sites with more than 80% of samples covered, we obtained information for 43,420 CpG sites using the *ASM*_*snp*_. Using the tuple score, we obtained summaries for 1,849,831 CpG pairs. Within the **DAMEfinder** pipeline, we generated multi-dimensional scaling (MDS) plots using each score (Figures 5A and B), and observed that both scores are able to recover distinct CRC phenotypes. However using the *ASM*_*tuple*_ score, samples cluster according to tissue type (normals, CIMP cancer and non-CIMP cancer) (Figure 5A), whereas using the *ASM*_*snp*_ score, only the two cancer types are distinguishable, while the normal tissues cluster with their matched cancers (Figure 5B).

**Figure 5.**
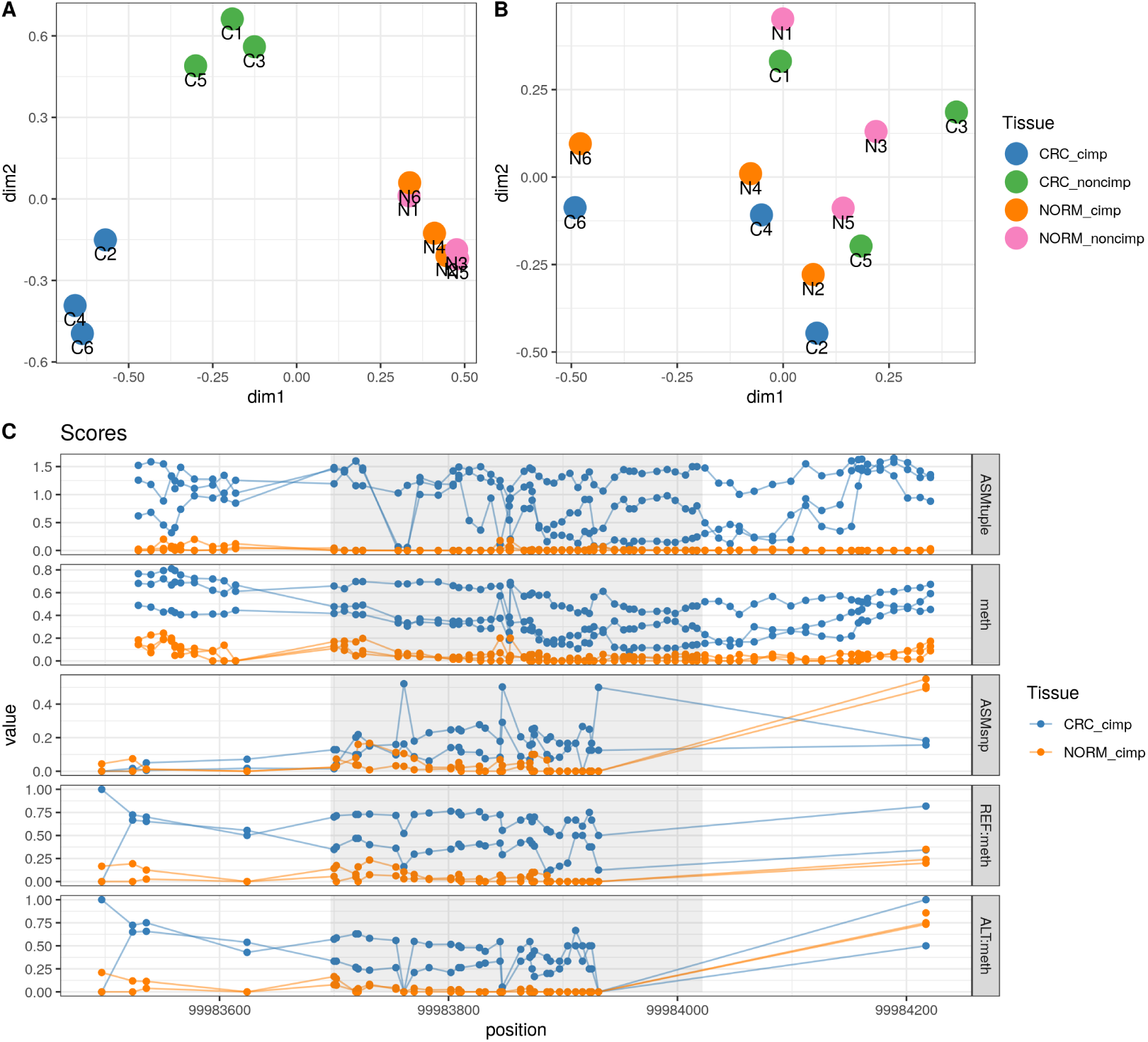
ASM scores on the CRC dataset. **A.** MDS plot of all the samples in the CRC dataset, based on all the the *ASM*_*tuple*_ scores. Scores were square-root transformed before plotting. **B.** MDS plot based on the *ASM*_*snp*_ scores. Scores were arcsine transformed. MDS plots were generated with the *plotMDS* function from **limma** and the top 1000 most variable positions. N: normal mucosa; C: CRC. Each pair of samples from the 6 patients with CRC are numbered from 1 to 6. **C.** A DAME detected in CIMP CRCs using the *ASM*_*tuple*_ score shows a higher signal than using the *ASM*_*snp*_ score. Region shown is located on chr9:99,983,697-99,984,022, shaded region in the center corresponds to the DAME. Tracks for methylation levels (meth) and methylation levels in reference and alternative alleles (based on SNP in chr9:99,983,812) is also shown. Points in *ASM*_*tuple*_ and meth tracks correspond to intermediate positions between a pair of CpG sites. Points in the rest of tracks correspond to CpG sites.

We performed DAME detection on each score independently using the Cluster-wise correction (Supplementary Figure 2, Additional File 1 for p-values of both Cluster-wise correction and Permutations). When using the *ASM*_*snp*_ score, we could not detect DAMEs with an adjusted p-value below 0.05. Using the *ASM*_*tuple*_ score, we were able to detect 4,051 DAMEs in the CIMP samples (versus matched normal samples), and 258 in the non-CIMP samples. We noticed that regions detected using *ASM*_*tuple*_ were also detected using *ASM*_*snp*_, but with lower strength of signal and with p-values above a cutoff of 0.05 (one example in Figure 5C), and other regions showing the contradicting changes in ASM (one example in Supplementary Figures 3-4, Additional File 1). Additionally, we found DAMEs corresponding to known regions exhibiting loss of imprinting in cancer, including those in the genes *MEG3, H19*, and *GNAS* [13, 50] (Figure 6).

**Figure 6.**
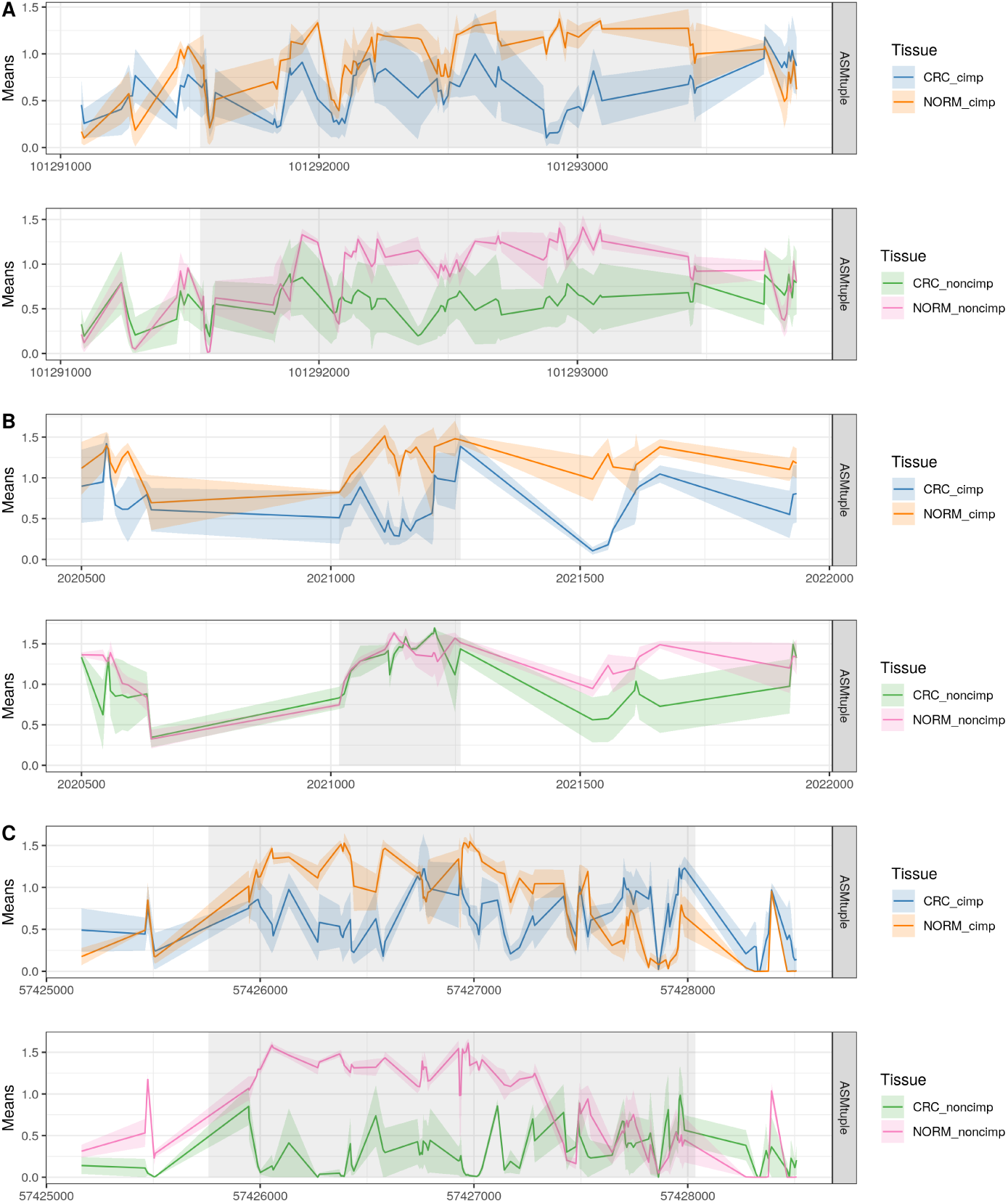
DAMEs overlapping known loci exhibiting loss of imprinting in colorectal cancer. **A.** DAME located in chr14:101,291,540-101,293,480, upstream the imprinted *MEG3* gene. The loss of imprinting was significant in both types of CRCs. **B.** DAME located in chr11:2,021,017-2,021,260, upstream the imprinted *H19* gene. Loss of imprinting only occurred in CIMP CRCs. **C.** DAME in the *GNAS* gene located in chr20:57,425,758-57,428,036. Loss of imprinting was detected in both types of CRCs. Y-axis in all panels corresponds to *ASM*_*tuple*_ means. Lines connect means at intermediate positions between a pair of CpG sites. Shared areas correspond to confidence intervals at each position (standard errors of the mean).

Considering the high number of DAMEs detected in the CIMP contrast compared to the non-CIMP contrast, we thought this could be a consequence of hypermethylation in CIMP [51], and a typical DMR (differentially methylated region) analysis would be able to detect these same regions. To corroborate this, we performed a DMR analysis on the CIMP and non-CIMP contrasts using the **dmrseq** R package [46] (Supplementary Figure 5, Additional File 1 for top DAMEs and DMRs per comparison). We found that from the 6,753 DMRs (5,040 hypermethylated, 1,713 hypomethylated) detected in the CIMP comparison, 2,285 overlap with DAMEs (hypermethylated DMRs = 32%, hypomethylated DMRs = 1.7% from total DMRs), and from 13,220 DMRs in the non-CIMP comparison, only 164 overlap (hypermethylated DMRs = 0.57%, hypomethylated DMRs = 0.66%) (Table 1).

**Table 1.**
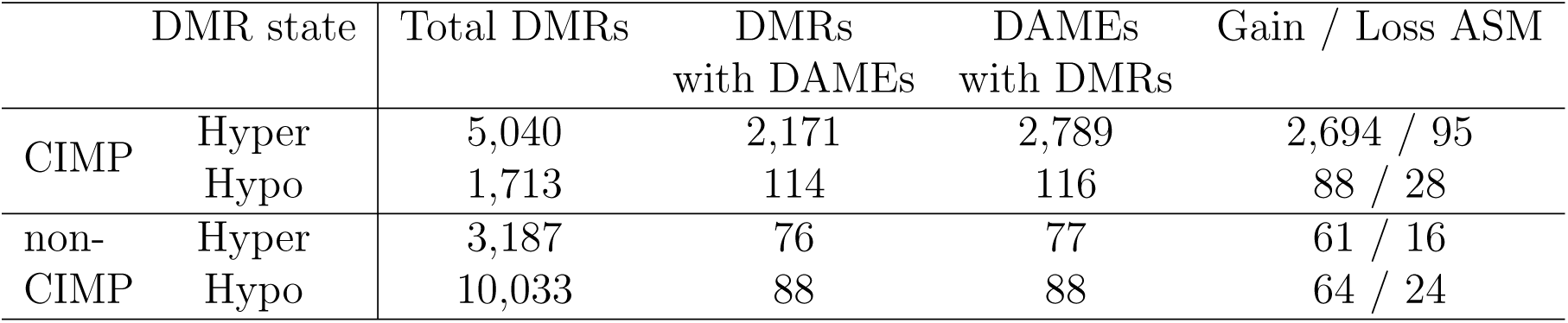
DMRs overlapping DAMEs. Hyper or hypo-methylated DMR refers to the increase or decrease of methylation in cancers in comparison with paired normal samples, while gain or loss of ASM refers to whether cancers have more or less allele-specificity than paired normal samples.

Because of this overlap, we conclude that a proportion (1,146 [28%] in CIMP, 93 [36%] in non-CIMP) of DAMEs would not be detected via a typical DMR analysis. Figure 7 shows 4 examples of DAMEs missed by the DMR detection. In principle, these regions exhibit differential methylation according to the global methylation levels (bottom panels of each region), however the hypermethylation reaches intermediate values, which might not represent a sufficiently high effect size to be detected. However, in the context of differential ASM, these intermediate values are highly scored, based also on the allele-specificity of the change. Therefore, even though these are not highly ranked DAMEs, they were still included as such.

**Figure 7.**
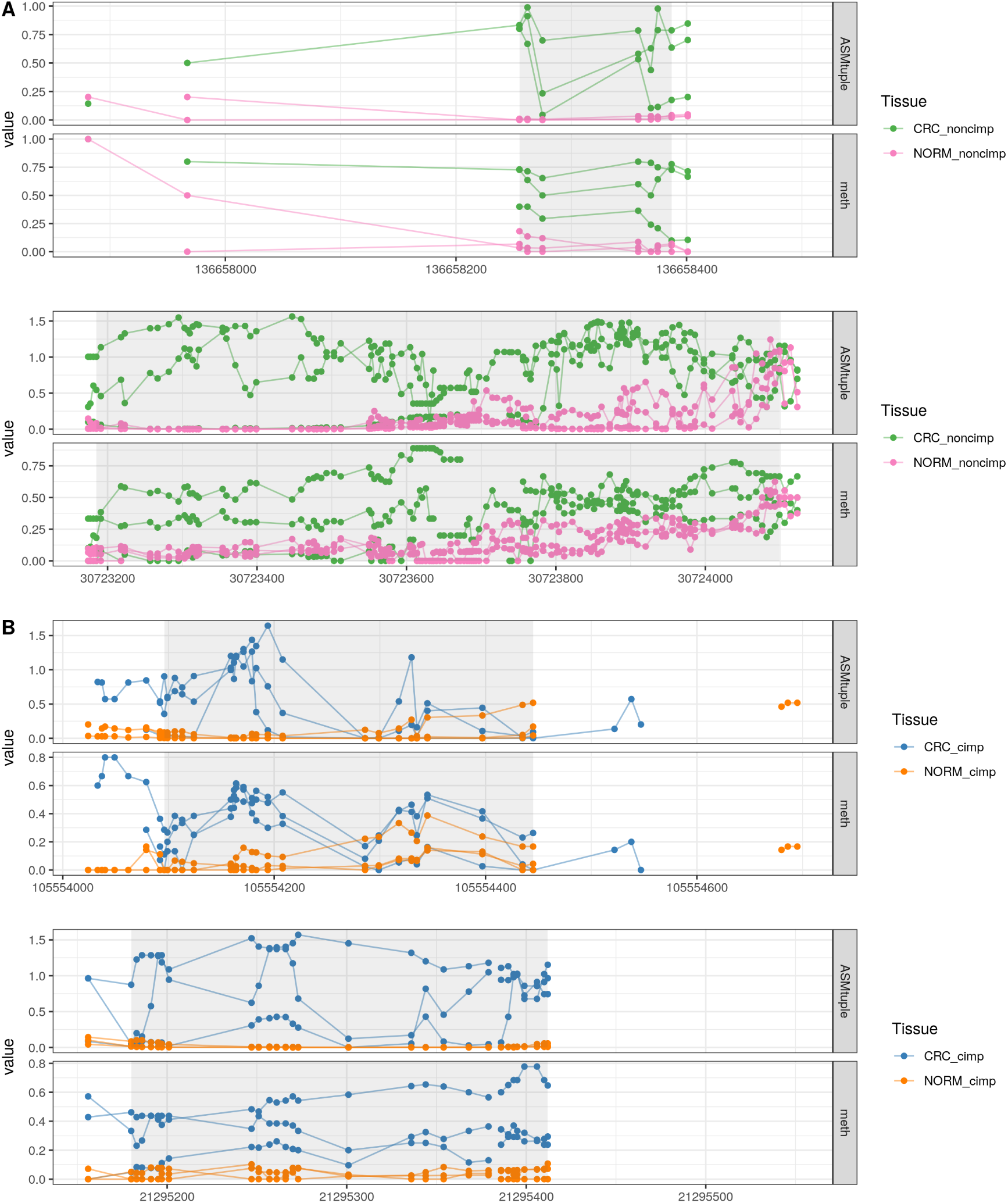
DAMEs not detected as DMRs. **A.** Two different DAMEs in non-CIMP, the first located in chr9:136,658,255-136,658,387, and the second located in chr4:30,723,185-30,724,099. **B.** Two different DAMEs in CIMP, the first in chr14:105,554,096-105,554,445; the second in chr16:21,295,180-21,295,412. Y-axis corresponds to *ASM*_*tuple*_ or methylation. Points correspond to intermediate positions between a pair of CpG sites.

## Discussion

We have developed a scoring method that provides a measure of allele-specific methylation, and developed a method (DAMEfinder) that detects regions that display loss or gain of allele-specific methylation, by leveraging existing methods into a single framework. We offer the possibility to detect regions exhibiting ASM based on genotype information (*ASM*_*snp*_), or independent from it (*ASM*_*tuple*_). The latter offers a novel approach for identifying different types of ASM, such as imprinted, non-imprinted, XCI, and new types yet to be described.

Compared to existing scores (**allelicmeth**, deviations from 50% methylation), *ASM*_*tuple*_ showed favourable performance at identifying individual cases of ASM at different coverage levels. The scaled methylation also demonstrated high sensitivity and specificity, and as the true ASM score (*ASM*_*snp*_) and coverage were increased, results were close to those of the *ASM*_*tuple*_ score. Nonetheless, the advantage of using the *ASM*_*tuple*_ score is the flexibility in its implementation; specifically, the weight that is added to the log-odd ratio can be adapted to the user’s needs. As an example, one could argue that a 50:50 proportion of methylated to unmethylated reads is not a good indicator of ASM. This assumption can be relaxed or changed within the model by changing the level of departure *ϵ* in the weight calculation.

In contrast, the **allelicmeth** score reduced its performance when the true ASM value was increased. As for **amrfinder**, we believe defining ASM as regional is a nice implementation in this method, and can make ASM interpretation and visualization easier. However, the definition of regions is done for each sample independently, and this does not allow for a direct comparison between samples. This is the main reason why our ASM scores are not regional. Our method focuses on obtaining regions of consistent *change* in ASM between conditions relative to the variability, which in turn implies consistent ASM in the majority of samples from an experimental condition.

Our *ASM*_*tuple*_ score was able to distinguish female from male samples based on XCI. When analyzing the entire genome however, we did not find differences between males and females. The fact that the entire female chromosome X does not contain high ASM, or that the global distribution of methylation is not skewed towards intermediate values has been shown before [52]. The presence of genes escaping XCI may also affect global ASM. It is known that 15% of genes escape XCI, and an additional 10% vary in the inactivation state among the female population [53]. Therefore, a mixture of ASM scores in females is an accurate reflection of the complex dynamics of XCI.

We were also able to validate the score by comparing the promoters of 89 known imprinted genes with the rest of the genome. We observed an increase in the ASM of imprinted genes, with a bimodal distribution of ASM scores. This can be a reflection of tissue or cell type specificity in imprinted genes, meaning not all known imprinted genes show ASM throughout the somatic cell lineage, as is traditionally assumed [54]. Studies have reported tissue and cell type-specific allelic expression [55, 56] and tissue-specific ASM [23] in known imprinted genes, supporting our finding that imprinting is not equally maintained in all genes in every tissue and/or cell type.

Another aspect that could easily affect the range of ASM scores is cell heterogeneity, where we may expect a mixture of methylated and unmethylated alleles. The fact that the ASM scores observed in both the CRC and Blood datasets are continuous is likely a reflection of this. We expect ASM to be an all or none phenomenon, where “real” ASM should be either fully allele-specific (one allele fully methylated and the other fully unmethylated) or not (either both fully methylated, or both fully unmethylated). Additionally, reads from the colorectal cancer dataset were sequenced from cancerous tissue, which is typically associated with high intra-tumor heterogeneity of several biological features, including cellular morphology and gene expression [57]. Our method does not account for this additional variability, and we recognize this as a limitation. However, we believe the ASM scores are still robust enough to detect allelic patterns as shown by the recovery of the colorectal cancer subtypes in Figure 5 and that even changes in cell composition, which would also affect DMR detection, can be interesting events to understand.

To obtain all-or-none ASM, single cell BS-seq (scBS-seq) data may become the most suitable high-throughput technology. Previous studies have shown the use of scBS-seq to detect heterogeneity within a single cell type [58] and cell states [59]. However, the accurate detection of methylation from scBS-seq is still a difficult task, mainly due to the extensive DNA damage from the bisulfite treatment. There are currently around 21 different protocols to profile single cell DNA methylation, mostly bisulfite-based, each one aiming at improving recovery of CpGs and mapping efficiency [60]. However, it has not been established how these methods compare to each other, and a consistent framework for their data analysis does not exist, as is the case for bulk BS-seq protocols. Therefore there is still work ahead to precisely quantify ASM using scBS-seq.

Regarding DAME detection, we offer two strategies that differ in the statistical stringency. In our experience, fewer regions are obtained by permuting the group labels, since the FDR control is more conservative. However, more regions can always be detected by setting the *K* threshold lower, while still controlling the FDR. The Cluster-wise correction, or Simes method is less conservative, and therefore can be used as an alternative to extract more detection power. This is likely because of the global hypothesis tested at each DAME, where at least one CpG site in a region is changed.

We applied DAMEfinder to a real dataset to detect DAMEs in CIMP and non-CIMP cancers (versus paired normal samples). We found that the *ASM*_*tuple*_ and *ASM*_*snp*_ scores are consistent in describing the CIMP status of samples, but as expected, the *ASM*_*snp*_ score was dominated by SD-ASM, because its calculation relies on the heterozygous SNPs of each sample; paired samples thus clustered with each other not by tissue, as observed with the *ASM*_*tuple*_ score. Additionally, *ASM*_*tuple*_ typically detected more DAMEs, which we attribute to two reasons. First, there are ∼40x more places in the genome where *ASM*_*tuple*_ can be calculated. Second, because the tuple score is a more general calculation, i.e., it quantifies the mixing of methylated and unmethylated reads, instead of relying on allele information.

We also compared the DAME detection to a typical DMR analysis of the same samples, and found that DMRs detected may or may not include DAMEs. Most DMRs overlapping DAMEs were hypermethylated in CIMP cancers, which led us to conclude that most DAMEs reflected gain of ASM from a low methylation baseline. This result shows how differential ASM is a more refined definition of differential methylation, and can therefore provide additional information regarding methylation disruptions in disease (or different conditions).

## Conclusion

Cytosine methylation restricted to only one allele, i.e., ASM, is a particular pattern of methylation that should be approached differently than the rest of the human methylome. We have designed DAMEfinder to screen for ASM and identify regions of differential ASM. The latter can be viewed as a special case of differential methylation. Previous studies have quantified ASM within one sample, however, to our knowledge, there is no method that identifies loss or gain of ASM between conditions. DAMEfinder fills this gap. Studying changes in ASM can help us understand epigenetic processes in development and diseases. To this aim, further studies are necessary to associate ASM to allele specific gene expression and to verify whether gain or loss of ASM would affect gene dosage and eventually phenotypes.

## Methods

The code used to generate the article figures and data processing is available from https://github.com/markrobinsonuzh/allele_specificity_paper. The R package is available from https://github.com/markrobinsonuzh/DAMEfinder.

### Data Sets

#### Colorectal cancer (CRC) data set

The CRC data set came from our published study [51] describing the progression of a methylation signature from pre-cancerous lesions to colorectal cancer tissue in two types of CRC. We used 12 samples from 6 patients with sporadic cancer (arrayexpress accession number: E-MTAB-6949, Table 2). For each sample, DNA from both CRC lesion and normal mucosa was bisulfite treated and sequenced according the Roche SeqCapEpi CpGiant protocol, where only DNA captured by probes was sequenced. We analyzed 12 files in total. For details on data generation refer to [51].

**Table 2.** Colorectal cancer sample characteristics. *Sample ID changed from arrayexpress. C: CRC; N: paired sample of normal mucosa; non-CIMP: the mismatch repair gene MLH1 normally expressed; CIMP: MLH1 silenced by promoter hypermethylation.

| Sample ID* | CIMP status | Sex | Number of mapped reads | Average coverage | Average coverage in probes |
| --- | --- | --- | --- | --- | --- |
| N1 |  |  | 76,801,310 | 3.025 | 78.06 |
| C1 | non-CIMP | F | 68,010,696 | 2.47 | 61.62 |
| N2 |  |  | 74,815,980 | 2.97 | 69.96 |
| C2 | CIMP | M | 62,122,636 | 2.47 | 63.16 |
| N3 |  |  | 66,608,688 | 2.64 | 63.88 |
| C3 | non-CIMP | M | 57,828,284 | 2.28 | 57.52 |
| N4 |  |  | 66,108,442 | 2.62 | 58.61 |
| C4 | CIMP | M | 59,390,888 | 2.35 | 61.25 |
| N5 |  |  | 70,070,214 | 2.56 | 59.0032 |
| C5 | non-CIMP | M | 68,575,884 | 2.50 | 49.98 |
| N6 |  |  | 59,056,548 | 2.15 | 49.52 |
| C6 | CIMP | F | 79,669,532 | 2.92 | 71.39 |

**Table 3.**
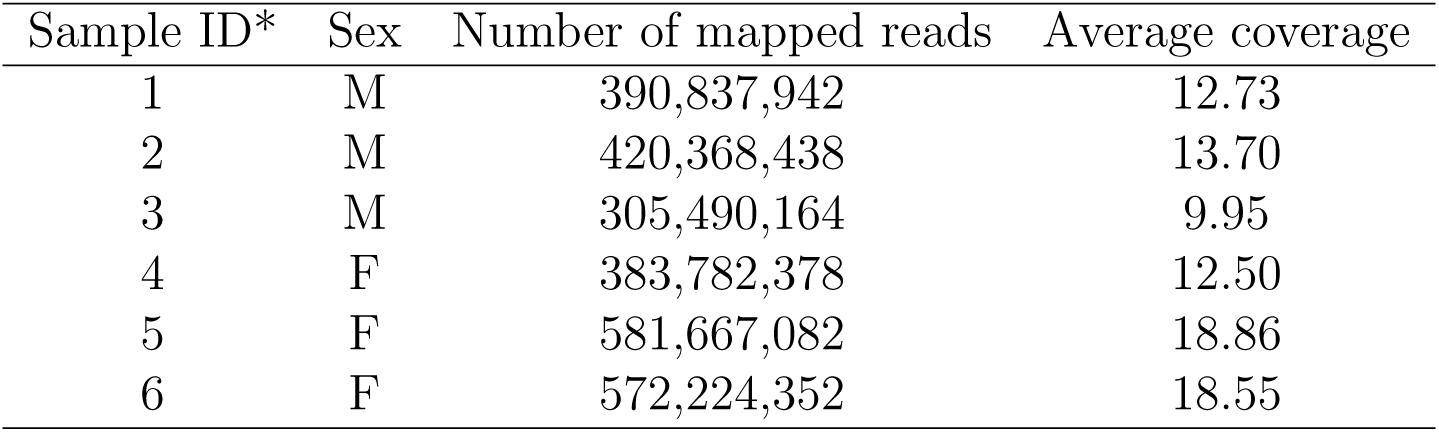
Blood data sample characteristics. *Sample ID changed from source.

#### Blood dataset

We used data generated by the Blueprint Consortium. We downloaded raw paired-end fastq files from venous blood of 3 healthy females and 3 healthy males (CD14-positive, CD16-negative classical monocyte, EGA dataset: EGAD00001002523).

### Quality control and mapping

Quality control was done using **fastQC** (version 0.11.4) [61]. The reads were subsequently trimmed using **TrimGalore!** (version 0.4.5) [62]. Reads were mapped to the reference genome using **bismark** (version 0.18.0). **Bowtie2** (version 2.2.9) was used to map to genome hg19 in the CRC data set, and hg38 in the Blood dataset. Duplicate reads were removed with the *deduplicate* command from **bismark**. Deduplicated bam files corresponding to technical replicates in the Blood data set were merged with **samtools merge** [63] **for each sample.**

### SNP calling

We extracted heterozygous SNPs from the CRC dataset bam files with **Bis-SNP** (version 1.0.0) [32] by running the *BisulfiteGenotyper* mode with default parameters, using the **dbSNP** (Build150) [64] generated VCF file from the NCBI Human Variation Sets (GRCh37p13, last modified:07-10-2017).

### methtuple

**Methtuple** (version 1.5.3) [40] was used to produce a list of unique tuples of size two and the corresponding MM, MU, UM, and UU counts where M stands for “methylated” and U for “unmethylated”. The bam files of each sample are those of PE reads and so they were sorted by queryname before using **methtuple**, as the tool demands it.

### tuple-based ASM Score

We used *γ*_1_ = *γ*_2_ = 0.5 and *ϵ* = 0.2 for all analyses, and allowed for a maximum distance of 150 base pairs between two CpGs in a tuple. Supplementary Figure 6, Additional File 1, show *ASM*_*tuple*_ diagnostic plots for the CRC dataset (and Supplementary Figure 7 with *ASM*_*snp*_).

#### *ASM*_*tuple*_ score transformation

We apply a square root transformation to the *ASM*_*tuple*_ score before running **limma**, to get a more stable mean-variance relationship.

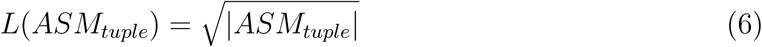

### allelicmeth

**allelicmeth** (**MethPipe** version 3.4.3) [33] is a tool that also detects ASM for a given sample directly from BS-seq reads. The tool is part of the **MethPipe** pipeline [65], which does not use standard bam files. We used commands from the pipeline to transform our **bismark** bam files from the CRC dataset into *mr* files, the input to **allelicmeth**. The output is a bed file with p-values for each pair of CpG sites, reflecting the degree of allele-specificity.

### amrfinder

**amrfinder** (**MethPipe** version 3.4.3) [33] also detects ASM from the BS-seq reads, however it generates regional scores. As with **allelicmeth**, we transformed **bismark** bam files from the CRC dataset into *mr* files, then ran *methstates* to generate *epiread* files, and used these to run **amrfinder** with default parameters. The output is a bed file with p-values for each genomic region with consistent ASM.

### Score evaluation

We converted the *ASM_snp_* into a tuple-*ASM_snp_* as 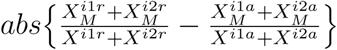, where 1 and 2 are the the first and second CpG site in a tuple *i*. We treated this converted score as true allele-specific methylation to test our scores at two thresholds: ≥ 0.5 and ≥ 0.8.

We transformed the p-values generated by **allelicmeth** and **amrfinder** with a negative log base 10. We assigned the same transformed p-values to all CpG tuples included in a single **amrfinder** region.

We also compared to a score based on whether the proportion of methylated reads to total number of reads deviates from 0.5, but transformed so a value of 0.5 is indicative of high ASM, and 1 or 0 is the lowest ASM. The score is 1 − 2(|*methylation* − 0.5|).

We used these four metrics to build ROC curves at different read coverages (5-9, 10-49 and ≥ 50) and at different thresholds of *ASM*_*snp*_, for a single normal mucosa sample in the CRC data set.

As an additional validation, we used the Blood dataset to obtain the *ASM*_*tuple*_ scores from the promoters of known imprinted genes reported in [11]. Only gene symbols that were traceable with **biomaRt** [66, 67] were included, and genes labelled to be imprinted in placenta were removed, as indicated in [68, 69].

### t-statistic calculation

From the **limma** R package [38], we use **lmFit** to fit a linear model for each CpG position, and **eBayes** to calculate a moderated t-statistic on the transformed *ASM*_*tuple*_ score, or on the *ASM*_*snp*_ score. For the former, we set the median of two CpGs in a tuple as the CpG position of that tuple. Transformed ASM scores across samples are given as input to **lmFit**, as well as a design matrix that specifies the conditions of the samples of interest. As specified in [38, 70], a CpG site-wise or tuple-wise linear model is defined as:

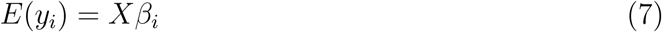

where for each CpG site or tuple *i*, we have a vector of ASM scores *y*_*i*_ and a design matrix *X* that relates these values to some coefficients of interest *β*_*i*_.

In the end, we test for a specific contrast that *H*_*o*_ : 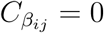.

### Smoothing

We group the positions into genomic clusters using the **clusterMaker** function from the **bumphunter** R package [39]. Then we use the **loessByCluster** function to perform loess within each cluster, and obtain 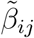, our smoothed estimate.

### FDR control evaluation

We selected 6 samples of normal tissue from the CRC dataset and calculated their *ASM*_*snp*_ scores as a baseline in the simulation. We divided the samples in 2 groups of 3. We generated 1038 clusters of CpGs with the **clusterMaker** function from the **bumphunter** package, and set a maximum distance between CpGs of 100 bp (Supplementary Figure 8, Additional File 1). We chose 20% of all clusters to be truly differential, and to each of them added effect to a number of randomly selected consecutive CpGs. The effect size is the same for every chosen CpG per cluster, and is obtained by inverse transform sampling of the form 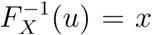, where *u ∼ Unif*(0.35, 0.75), and *F*_*X*_(*x*) the CDF of *Beta*(1, 2.5) [46] (Supplementary Figure 9, Additional File 1). Additionally, for each truly differential cluster, we randomly selected the sign of the effect size (positive or negative), as well as the group of samples that contains the effect size.

We generated 50 of these simulations, and for each of them, ran DAMEfinder with the cluster-wise correction, and the permutation correction (Supplementary Figure 10, Additional File 1 for distributions of null and observed areas) with three different *K* thresholds: 0.2, 0.5, 0.8. We used the **iCOBRA** R package (version 1.12.1) [71] to calculate TPR and FDR at different adjusted p-value thresholds: 0.01, 0.05, 0.1.

### DMR detection

We identified DMRs with the **dmrseq** R package (version 1.5.11) [46] for each cancer subtype. We specified the tissue via the *testCovariate* parameter (CIMP, non-CIMP or normal), and the patient with the *adjustCovariate* parameter. The *cutoff* parameter (cutoff of the single CpG coefficient that is used to discover candidate regions) was set as 0.05 and the rest of parameters were set as default.

## Author’s contributions

MM and MDR conceived the study. SO, DM and MDR wrote package, performed analyses. SO, GM and MDR wrote the paper. All authors read and approved the final manuscript.

## Funding

GM and SO acknowledge funding from the SNF grant 310030-160163/1. MDR acknowledges support from the University Research Priority Program Evolution in Action at the University of Zurich.

## Acknowledgements

The authors thank Abdullah Kahraman for technical support at previous stages of the project; Pierre-Luc Germain and Izaskun Mallona for feedback on the manuscript; and the Robinson lab for feedback on figures and analysis.

## Notes

#### Summary of Updates

Change "paternal" to "parental" in abstract; thanks to at_shian_su

## References

[1] Cedar, H. and Bergman, Y. (2009). Linking DNA methylation and histone modification: patterns and paradigms. Nature Reviews Genetics, 10:295–304.

[2] Bonasio, R., Tu, S., and Reinberg, D. (2010). Molecular signals of epigenetic states. Science, 330(6004):612–616.

[3] Bird, A. P. (1978). Use of restriction enzymes to study eukaryotic DNA methylation: II. the symmetry of methylated sites supports semi-conservative copying of the methylation pattern. Journal of Molecular Biology, 118(1):49–60.

[4] Suzuki, M. M. and Bird, A. (2008). DNA methylation landscapes: provocative insights from epigenomics. Nature Reviews Genetics, 9:465–476.

[5] Bergman, Y. and Cedar, H. (2013). DNA methylation dynamics in health and disease. Nature Structural & Molecular Biology, 20:274–281.

[6] Clark, S. J., Statham, A., Stirzaker, C., Molloy, P. L., and Frommer, M. (2006). DNA methylation: bisulphite modification and analysis. Nature protocols, 1(5):2353–2364.

[7] Liu, Y., Siejka-Zielinska, P., Velikova, G., Bi, Y., Yuan, F., Tomkova, M., et al. (2019). Bisulfite-free direct detection of 5-methylcytosine and 5-hydroxymethylcytosine at base resolution. Nature Biotechnology, 37(4):424–429.

[8] Reddington, J. P., Pennings, S., and Meehan, R. R. (2013). Non-canonical functions of the DNA methylome in gene regulation. Biochemical Journal, 451(1):13–23.

[9] Ferguson-Smith, A. C. (2011). Genomic imprinting: the emergence of an epigenetic paradigm. Nature Reviews Genetics, 12:565–575.

[10] Bartolomei, M. S. and Ferguson-Smith, A. C. (2011). Mammalian genomic imprinting. Cold Spring Harbor Perspectives in Biology, 3(7).

[11] Tucci, V., Isles, A. R., Kelsey, G., Ferguson-Smith, A. C., and the Erice Imprinting Group (2019). Genomic imprinting and physiological processes in mammals. Cell, 176(5):952–965.

[12] Knoll, J. H. M., Nicholls, R. D., Magenis, R. E., Graham Jr., J. M., Lalande, M., Latt, S. A., et al. (1989). Angelman and prader-willi syndromes share a common chromosome 15 deletion but differ in parental origin of the deletion. American Journal of Medical Genetics, 32(2):285–290.

[13] Cui, H., Onyango, P., Brandenburg, S., Wu, Y., Hsieh, C.-L., and Feinberg, A. P. (2002). Loss of imprinting in colorectal cancer linked to hypomethylation of H19 and IGF2. Cancer Research, 62(22):6442–6446.

[14] Schwienbacher, C., Gramantieri, L., Scelfo, R., Veronese, A., Calin, G. A., Bolondi, L., et al. (2000). Gain of imprinting at chromosome 11p15: A pathogenetic mechanism identified in human hepatocarcinomas. Proceedings of the National Academy of Sciences, 97(10):5445–5449.

[15] Lyon, M. F. (1961). Gene action in the X-chromosome of the Mouse (Mus musculus L.). Nature, 190(4773):372–373.

[16] Kerkel, K., Spadola, A., Yuan, E., Kosek, J., Jiang, L., Hod, E., et al. (2008). Genomic surveys by methylation-sensitive SNP analysis identify sequence-dependent allele-specific DNA methylation. Nature Genetics, 40:904–908.

[17] Schalkwyk, L. C., Meaburn, E. L., Smith, R., Dempster, E. L., Jeffries, A. R., Davies, M. N., et al. (2010). Allelic skewing of DNA methylation is widespread across the genome. The American Journal of Human Genetics, 86(2):196–212.

[18] Tycko, B. (2010). Allele-specific DNA methylation: beyond imprinting. Human Molecular Genetics, 19(R2):R210–R220.

[19] Gertz, J., Varley, K. E., Reddy, T. E., Bowling, K. M., Pauli, F., Parker, S. L., et al. (2011). Analysis of DNA methylation in a three-generation family reveals widespread genetic influence on epigenetic regulation. PLOS Genetics, 7(8):1–10.

[20] Onuchic, V., Lurie, E., Carrero, I., Pawliczek, P., Patel, R. Y., Rozowsky, J., et al. (2018). Allele-specific epigenome maps reveal sequence-dependent stochastic switching at regulatory loci. Science, 361(6409).

[21] Wang, H., Lou, D., and Wang, Z. (2019). Crosstalk of genetic variants, allele-specific DNA methylation, and environmental factors for complex disease risk. Frontiers in Genetics, 9:695.

[22] Do, C., Lang, C. F., Lin, J., Darbary, H., Krupska, I., Gaba, A., et al. (2016). Mechanisms and disease associations of haplotype-dependent allele-specific DNA methylation. The American Journal of Human Genetics, 98(5):934–955.

[23] Marzi, S. J., Meaburn, E. L., Dempster, E. L., Lunnon, K., Paya-Cano, J. L., Smith, R. G., et al. (2016). Tissue-specific patterns of allelically-skewed DNA methylation. Epigenetics, 11(1):24–35.

[24] Faraday, N., Yanek, L. R., Yang, X. P., Mathias, R., Herrera-Galeano, J. E., Suktitipat, B., et al. (2011). Identification of a specific intronic PEAR1 gene variant associated with greater platelet aggregability and protein expression. Blood, 118(12):3367–3375.

[25] Klengel, T., Mehta, D., Anacker, C., Rex-Haffner, M., Pruessner, J. C., Pariante, C. M., et al. (2012). Allele-specific FKBP5 DNA demethylation mediates gene-childhood trauma interactions. Nature Neuroscience, 16:33–41.

[26] Zhang, Y., Rohde, C., Reinhardt, R., Voelcker-Rehage, C., and Jeltsch, A. (2009). Non-imprinted allele-specific DNA methylation on human autosomes. Genome Biology, 10(12):R138.

[27] Serre, D., Lee, B. H., and Ting, A. H. (2009). MBD-isolated Genome Sequencing provides a high-throughput and comprehensive survey of DNA methylation in the human genome. Nucleic Acids Research, 38(2):391–399.

[28] Down, T. A., Rakyan, V. K., Turner, D. J., Flicek, P., Li, H., Kulesha, E., et al. (2008). A bayesian deconvolution strategy for immunoprecipitation-based DNA methylome analysis. Nature Biotechnology, 26:779–785.

[29] Laird, P. W. (2010). Principles and challenges of genome-wide DNA methylation analysis. Nature Reviews Genetics, 11:191–203.

[30] Cheung, W. A., Shao, X., Morin, A., Siroux, V., Kwan, T., Ge, B., et al. (2017). Functional variation in allelic methylomes underscores a strong genetic contribution and reveals novel epigenetic alterations in the human epigenome. Genome Biology, 18(1):50.

[31] Zhu, P., Guo, H., Ren, Y., Hou, Y., Dong, J., Li, R., et al. (2018). Single-cell DNA methylome sequencing of human preimplantation embryos. Nature Genetics, 50(1):12–19.

[32] Liu, Y., Siegmund, K. D., Laird, P. W., and Berman, B. P. (2012). Bis-SNP: Combined DNA methylation and SNP calling for Bisulfite-seq data. Genome Biology, 13(7):R61.

[33] Fang, F., Hodges, E., Molaro, A., Dean, M., Hannon, G. J., and Smith, A. D. (2012). Genomic landscape of human allele-specific DNA methylation. Proceedings of the National Academy of Sciences of the United States of America, 109(19):7332–7.

[34] Martos, S. N., Li, T., Ramos, R. B., Lou, D., Dai, H., Xu, J.-C., et al. (2017). Two approaches reveal a new paradigm of ‘switchable or genetics-influenced allele-specific DNA methylation’ with potential in human disease. Cell Discovery, 3:17038.

[35] Robinson, M. D., Kahraman, A., Law, C. W., Lindsay, H., Nowicka, M., Weber, L. M., and Zhou, X. (2014). Statistical methods for detecting differentially methylated loci and regions. Frontiers in Genetics, 5:324.

[36] Shafi, A., Mitrea, C., Nguyen, T., and Draghici, S. (2018). A survey of the approaches for identifying differential methylation using bisulfite sequencing data. Briefings in Bioinformatics, 19:737–753.

[37] R Core Team (2019). R: A Language and Environment for Statistical Computing. R Foundation for Statistical Computing, Vienna, Austria.

[38] Ritchie, M. E., Phipson, B., Wu, D., Hu, Y., Law, C. W., Shi, W., and Smyth, G. K. (2015). limma powers differential expression analyses for RNA-sequencing and microarray studies. Nucleic Acids Research, 43(7):e47.

[39] Jaffe, A. E., Murakami, P., Lee, H., Leek, J. T., Fallin, M. D., Feinberg, A. P., and Irizarry, R. A. (2012). Bump hunting to identify differentially methylated regions in epigenetic epidemiology studies. International Journal of Epidemiology, 41(1):200–9.

[40] Hickey, P. (2014). Methtuple.

[41] Simes, R. J. (1986). An improved Bonferroni procedure for multiple tests of significance. Biometrika, 73(3):751–754.

[42] Krueger, F. and Andrews, S. R. (2011). Bismark: a flexible aligner and methylation caller for Bisulfite-Seq applications. Bioinformatics, 27(11):1571–1572.

[43] Lawrence, M., Huber, W., Pagès, H., Aboyoun, P., Carlson, M., Gentleman, R., et al. (2013). Software for computing and annotating genomic ranges. PLOS Computational Biology, 9(8):1–10.

[44] Shoemaker, R., Deng, J., Wang, W., and Zhang, K. (2010). Allele-specific methylation is prevalent and is contributed by cpg-snps in the human genome. Genome Research, 20(7):883–889.

[45] Affinito, O., Palumbo, D., Fierro, A., Cuomo, M., Riso, G. D., Monticelli, A., et al. (2019). Nucleotide distance influences co-methylation between nearby cpg sites. Genomics.

[46] Korthauer, K., Chakraborty, S., Benjamini, Y., and Irizarry, R. A. (2018). Detection and accurate false discovery rate control of differentially methylated regions from whole genome bisulfite sequencing. Biostatistics, page kxy007.

[47] Benjamini, Y. and Hochberg, Y. (1995). Controlling the false discovery rate: A practical and powerful approach to multiple testing. Journal of the Royal Statistical Society. Series B (Methodological), 57(1):289–300.

[48] Benjamini, Y. and Heller, R. (2008). Screening for partial conjunction hypotheses. Biometrics, 64(4):1215–1222.

[49] Lun, A. T. L. and Smyth, G. K. (2014). De novo detection of differentially bound regions for ChIP-seq data using peaks and windows: controlling error rates correctly. Nucleic Acids Research, 42(11):e95.

[50] Menigatti, M., Staiano, T., Manser, C. N., Bauerfeind, P., Komljenovic, A., Robinson, M. D., et al. (2013). Epigenetic silencing of monoallelically methylated miRNA loci in precancerous colorectal lesions. Oncogenesis, 2:e56.

[51] Parker, H. R., Orjuela, S., Oliveira, A. M., Cereatti, F., Sauter, M., Heinrich, H., et al. (2018). The proto CpG island methylator phenotype of sessile serrated adenomas/polyps. Epigenetics, 13(10-11):1088–1105.

[52] Duncan, C. G., Grimm, S. A., Morgan, D. L., Bushel, P. R., Bennett, B. D., Barnabas, B. B., et al. (2018). Dosage compensation and DNA methylation landscape of the X chromosome in mouse liver. Scientific Reports, 8(1):10138.

[53] Carrel, L. and Willard, H. F. (2005). X-inactivation profile reveals extensive variability in X-linked gene expression in females. Nature, 434(7031):400–404.

[54] Wood, A. J. and Oakey, R. J. (2006). Genomic imprinting in mammals: Emerging themes and established theories. PLOS Genetics, 2(11):1–9.

[55] Hippenmeyer, S., Johnson, R. L., and Luo, L. (2013). Mosaic analysis with double markers reveals cell-type-specific paternal growth dominance. Cell Reports, 3(3):960–967.

[56] Baran, Y., Subramaniam, M., Biton, A., Tukiainen, T., Tsang, E. K., Rivas, M. A., et al. (2015). The landscape of genomic imprinting across diverse adult human tissues. Genome Research, 25(7):927–936.

[57] Marusyk, A. and Polyak, K. (2010). Tumor heterogeneity: Causes and consequences. Biochimica et Biophysica Acta (BBA) - Reviews on Cancer, 1805(1):105–117.

[58] Smallwood, S. A., Lee, H. J., Angermueller, C., Krueger, F., Saadeh, H., Peat, J., et al. (2014). Single-cell genome-wide bisulfite sequencing for assessing epigenetic heterogeneity. Nature Methods, 11:817–820.

[59] Farlik, M., Sheffield, N., Nuzzo, A., Datlinger, P., Schönegger, A., Klughammer, J., et al. (2015). Single-cell DNA methylome sequencing and bioinformatic inference of epigenomic cell-state dynamics. Cell Reports, 10(8):1386 – 1397.

[60] Karemaker, I. D. and Vermeulen, M. (2018). Single-cell DNA methylation profiling: Technologies and biological applications. Trends in Biotechnology, 36(9):952 – 965.

[61] Andrews, S. (2015). fastqc.

[62] Krueger, F. (2017). Trim Galore!

[63] Li, H., Handsaker, B., Wysoker, A., Fennell, T., Ruan, J., Homer, N., et al. (2009). The Sequence Alignment/Map format and SAMtools. Bioinformatics, 25(16):2078–2079.

[64] Sherry, S. T., Ward, M.-H., Kholodov, M., Baker, J., Phan, L., Smigielski, E. M., and Sirotkin, K. (2001). dbSNP: the NCBI database of genetic variation. Nucleic Acids Research, 29(1):308–311.

[65] Song, Q., Decato, B., Hong, E. E., Zhou, M., Fang, F., Qu, J., et al. (2013). A reference methylome database and analysis pipeline to facilitate integrative and comparative epigenomics. PLOS ONE, 8(12):1–9.

[66] Durinck, S., Moreau, Y., Kasprzyk, A., Davis, S., De Moor, B., Brazma, A., and Huber, W. (2005). BioMart and Bioconductor: a powerful link between biological databases and microarray data analysis. Bioinformatics, 21:3439–3440.

[67] Durinck, S., Spellman, P. T., Birney, E., and Huber, W. (2009). Mapping identifiers for the integration of genomic datasets with the R/Bioconductor package biomaRt. Nature Protocols, 4:1184–1191.

[68] Court, F., Tayama, C., Romanelli, V., Martin-Trujillo, A., Iglesias-Platas, I., Okamura, K., et al. (2014). Genome-wide parent-of-origin DNA methylation analysis reveals the intricacies of human imprinting and suggests a germline methylation-independent mechanism of establishment. Genome Research, 24(4):554–569.

[69] Pervjakova, N., Kasela, S., Morris, A. P., Kals, M., Metspalu, A., Lindgren, C. M., et al. (2016). Imprinted genes and imprinting control regions show predominant intermediate methylation in adult somatic tissues. Epigenomics, 8(6):789–799.

[70] Smyth, G. K. (2004). Linear models and empirical bayes methods for assessing differential expression in microarray experiments. Statistical Applications in Genetics and Molecular Biology, 3(1):1–25.

[71] Soneson, C. and Robinson, M. D. (2016). iCOBRA: open, reproducible, standardized and live method benchmarking. Nature Methods, 13:283.

