## Additional File 1 for "DAMEfinder: A method to detect differential allele-specific methylation"

October 9, 2019

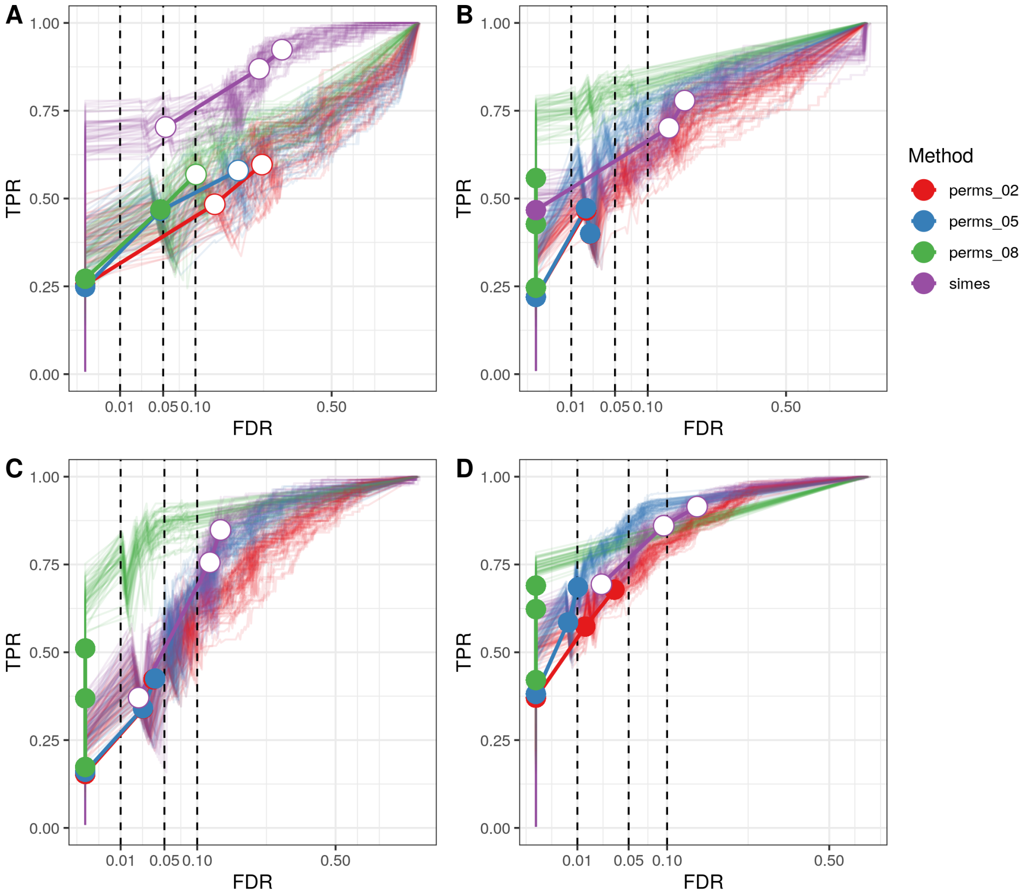


Supplementary Figure 1: Simulation runs for DAME detection. Same as Figure 3 in main text, with alternative parameters in simulation and detection. **A**. Maximum gap between CpG positions of 20 bps in simulated DAMEs. **B**. Maximum gap between CpG positions of 1000 bps in simulated DAMEs. **C**. Detection assuming prior variance is not constant (from eBayes). By default the function does not assume this. **D** Proportion of truly differential regions is 0.5. In main text the proportion is 0.2.

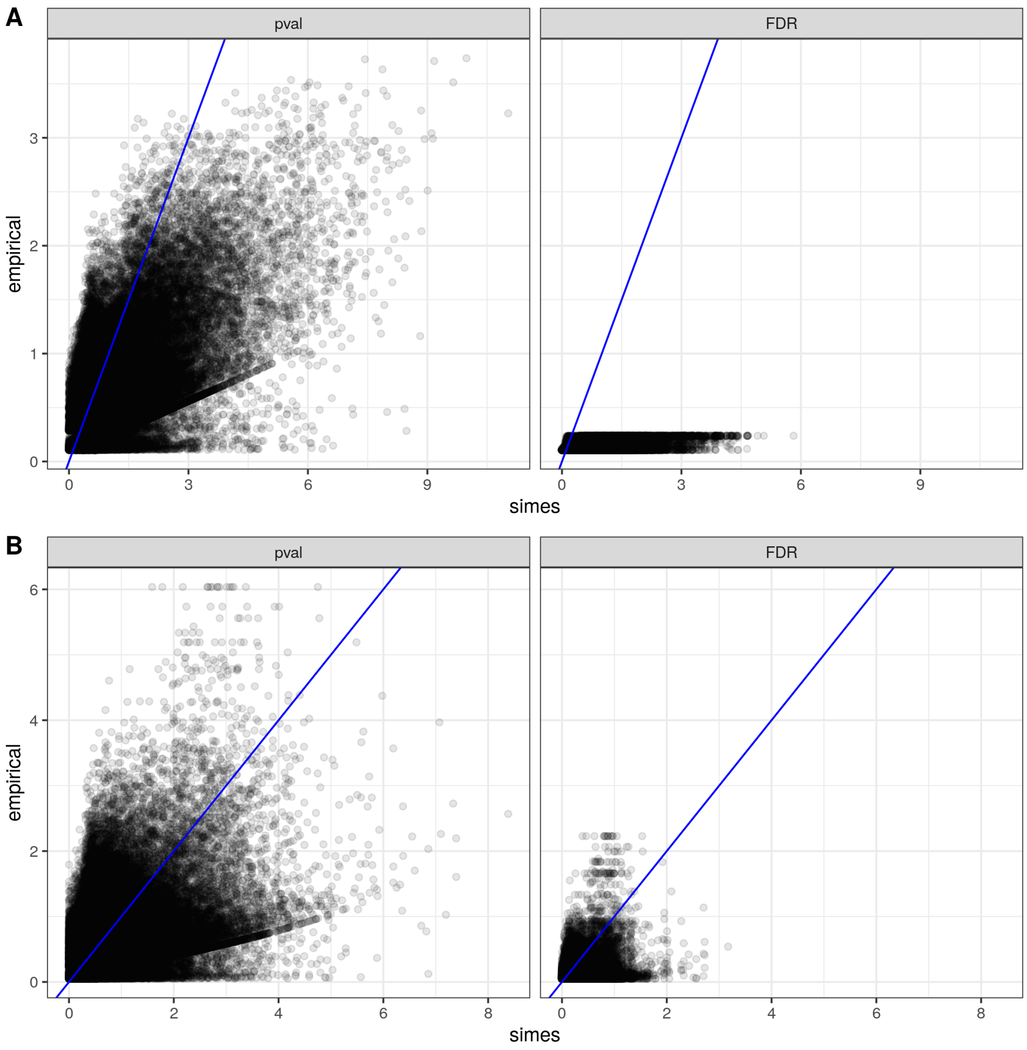


Supplementary Figure 2: Permutation and simes p-values assigned to CIMP (A) and non-CIMP (B) DAMEs. Left panels: p-values assigned to DAMEs using the permutation (empirical) method in the y-axis, and the cluster-wise correction (simes) in the x-axis. Right panels: Adjusted p-values using the Benjamini and Hochberg method. All axes are negative-log transformed.

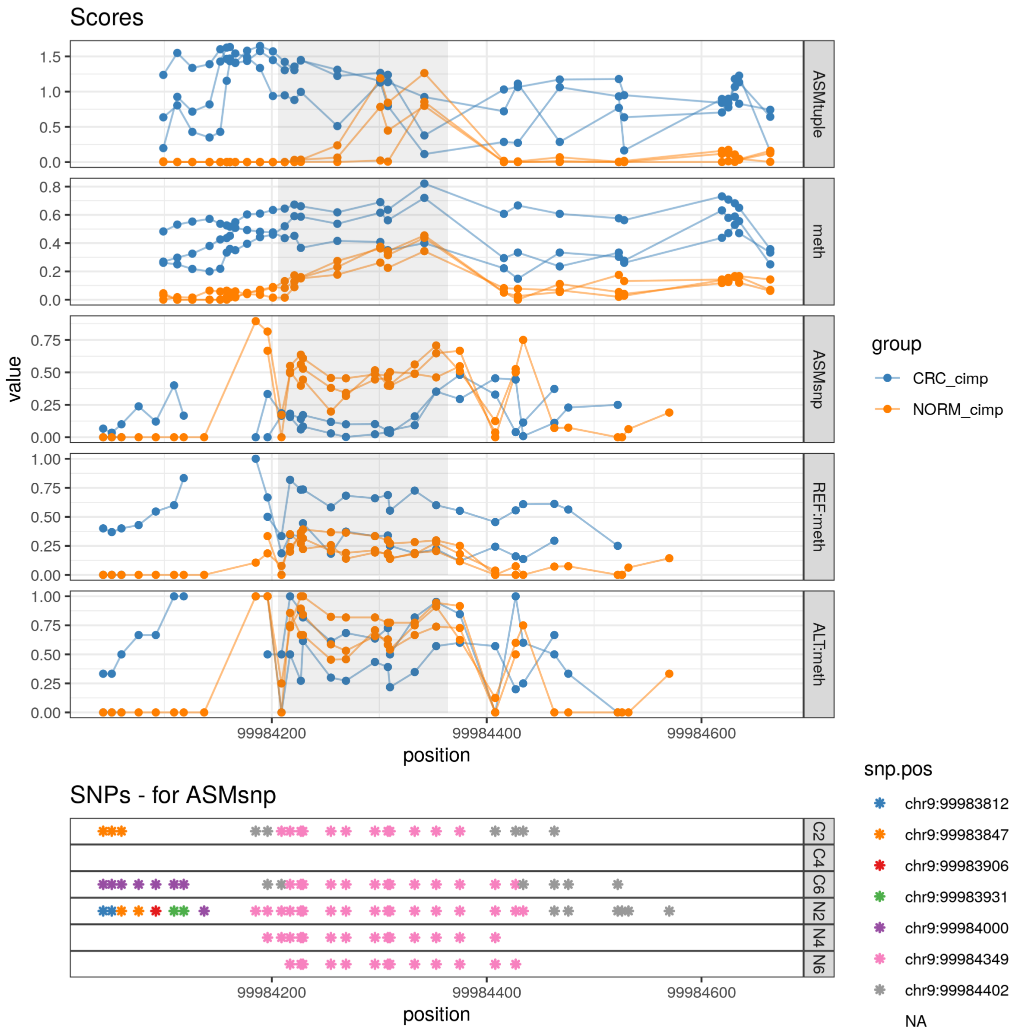


Supplementary Figure 3: Increase in $ASM_{tuple}$ score, decrease in $ASM_{snp}$ score. DAME located in chr9:99,984,206-99,984,364. $ASM_{tuple}$ shows gain of ASM in CRC samples, whereas $ASM_{snp}$ track shows loss of ASM in CRC samples.

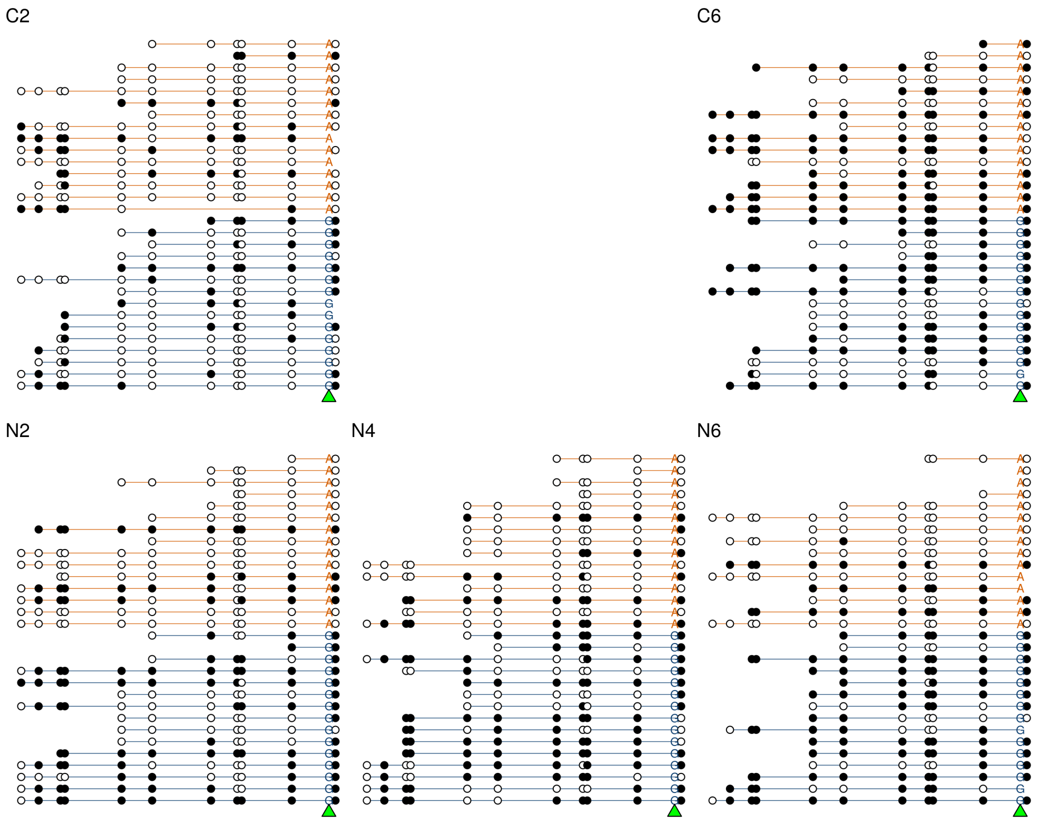


Supplementary Figure 4: Reads of DAME located in chr9:99,984,206-99,984,364 (from figure above 3) Reads sorted by allele show a different pattern than looking at unsorted reads (ignoring the allele). Figures labeled from C2 to N6 show the reads (horizontal lines colored by allele) overlapping the SNP (green triangle pointing to alleles) used to calculate this DAME from the $ASM_{snp}$ score. Samples with C are CRCs, samples with N are normal tissue. White circles are unmethylated CpG sites, black circles are methylated CpG sites. A random sample of 15 reads is shown for each allele.

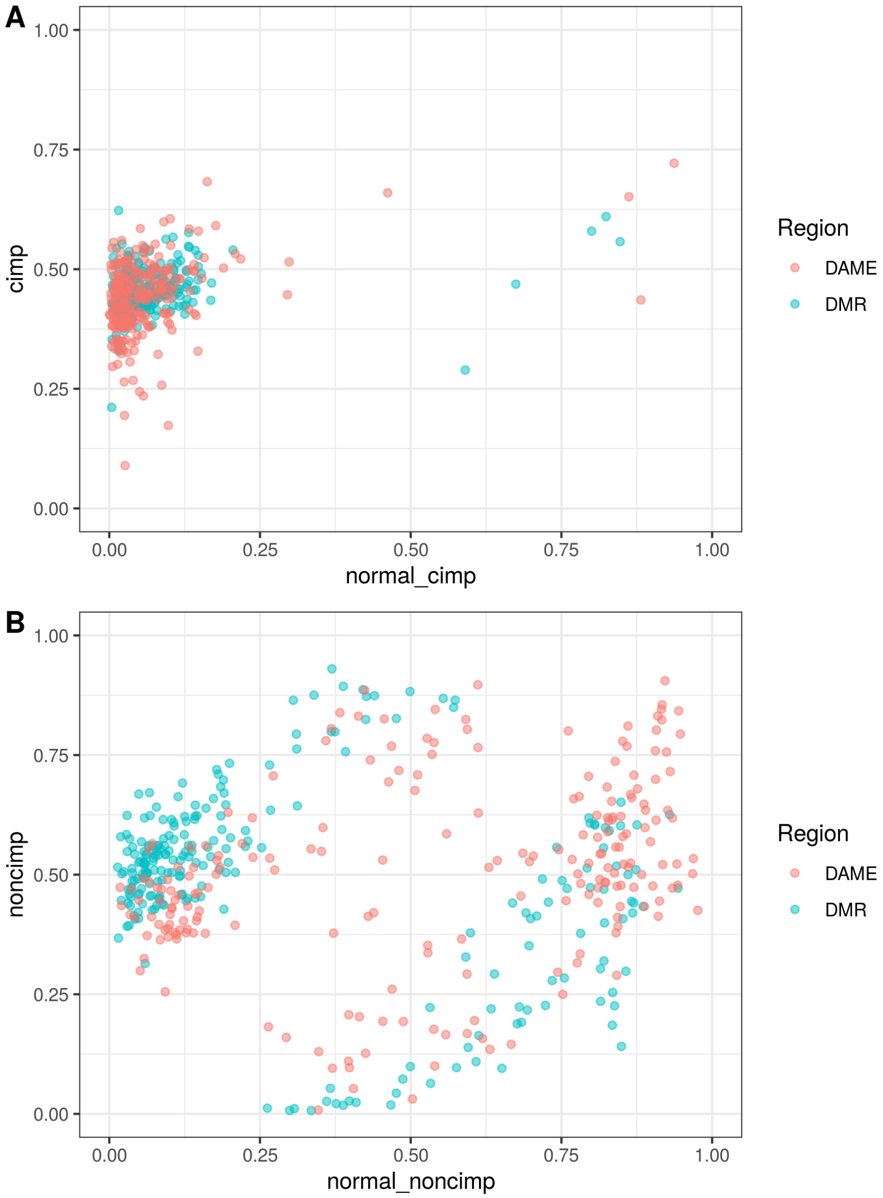


Supplementary Figure 5: Average methylation in top 250 DAMEs and DMRs. x-axis: Average methylation in normal tissue. y-axis: Average methylation in CRC tissue (average is across samples per group and then per region). **A**. CIMP CRCs **B**. non-CIMP CRCs.

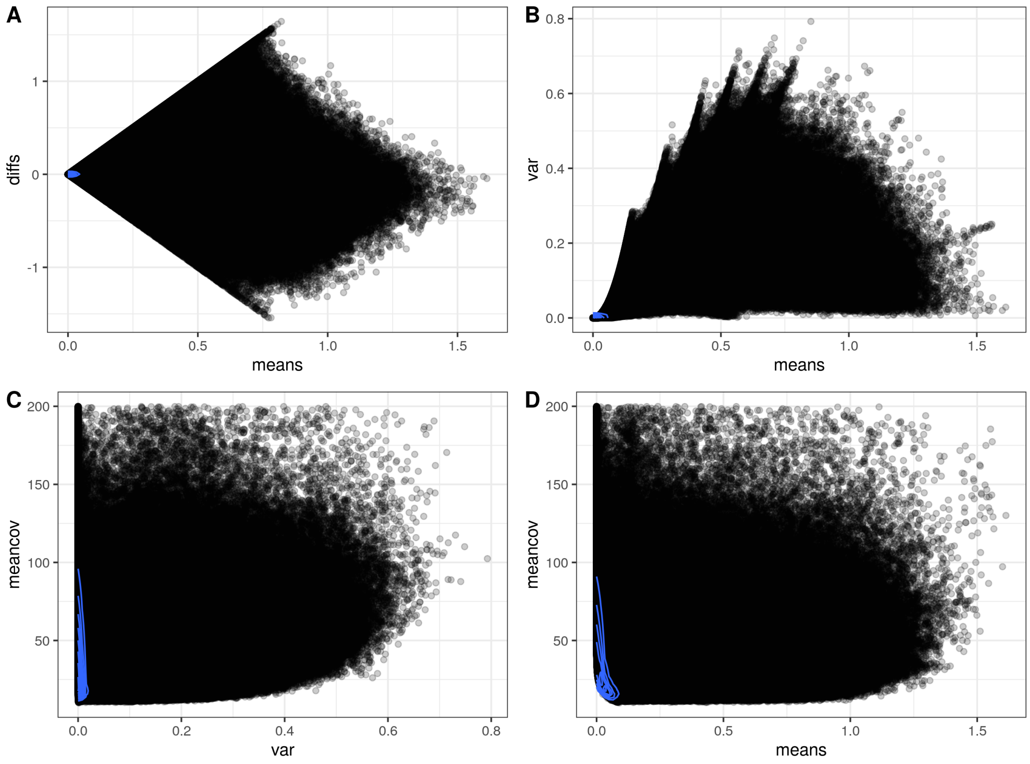


Supplementary Figure 6: $ASM_{tuple}$ score diagnostics in the CRC dataset. **A**. mean-difference plot, differences between all normals and all CRCs. **B**. mean-variance plot. **C**. Variance (x-axis) and mean coverage (y-axis). **D**. Mean of score (x-axis) and mean coverage (y-axis). Variance and mean do not depend on coverage. Each point corresponds to a CpG tuple.

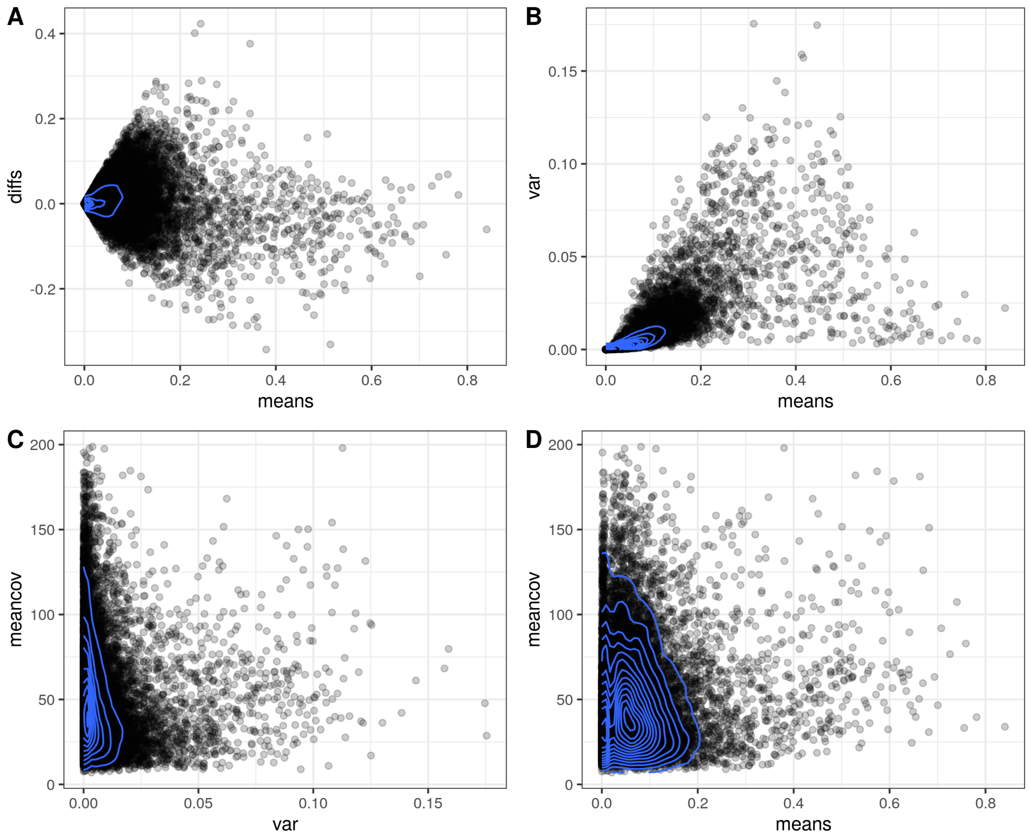


Supplementary Figure 7: $ASM_{snp}$ score diagnostics in the CRC dataset. **A**. Mean-difference plot, differences between all normals and all CRCs. **B**. Mean-variance plot. **C**. Variance (x-axis) and mean coverage (y-axis). **D**. Mean (x-axis) and mean coverage (y-axis). Each point corresponds to a CpG site.

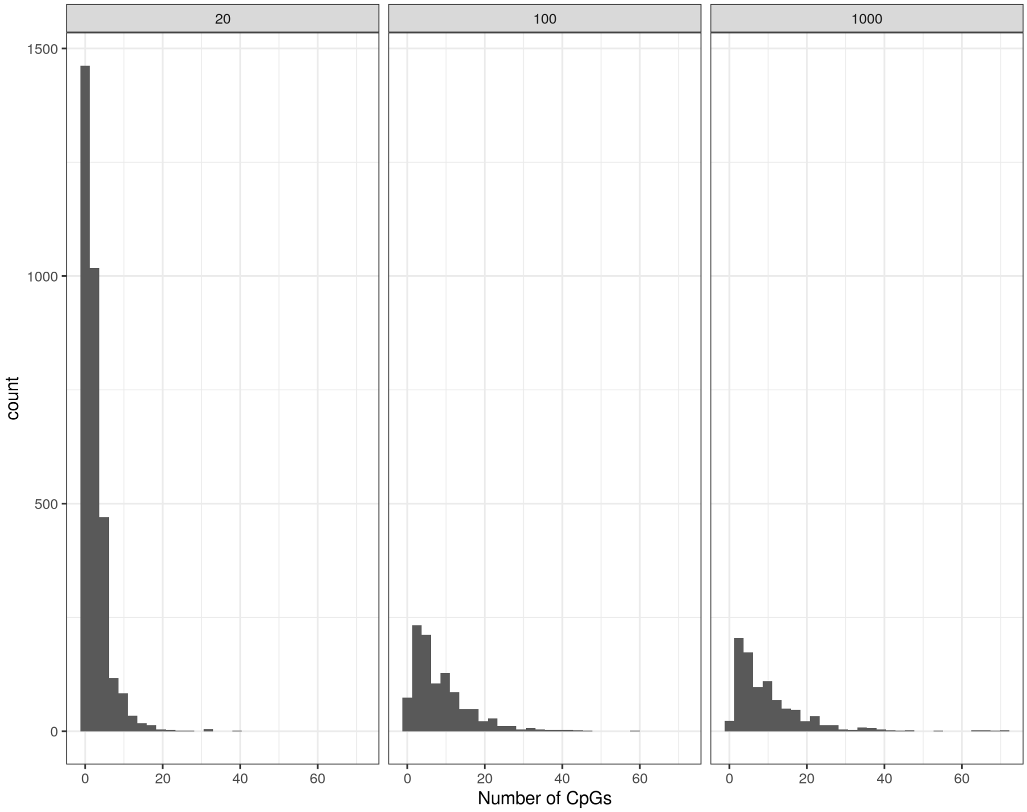


Supplementary Figure 8: Sizes of generated clusters (regions) for simulation. x-axis is the number of CpG sites per generated cluster. Simulation in main text was based on maxGap = 100 (middle facet). Other values for maxGap were used for Supp. Fig. 1

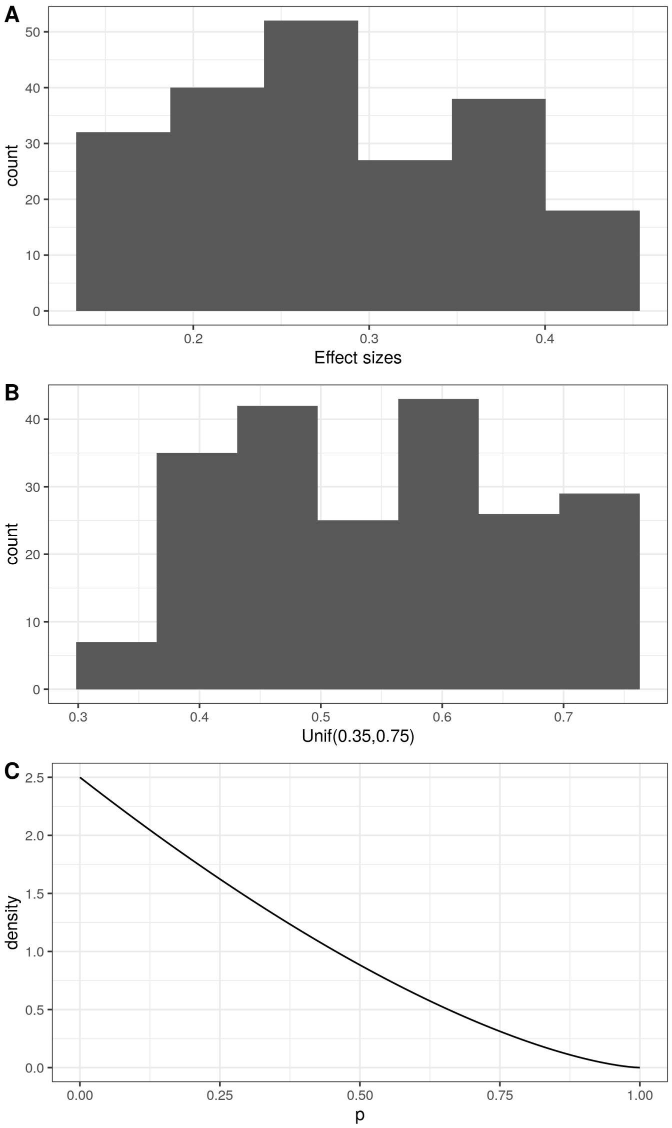


Supplementary Figure 9: Simulation parameters for effect size. **A**. Real effect sizes generated for 1 of the 50 simulations with inverse transform sampling of the form $F_X^{-1}(u) = x$, where $u \sim Unif(0.35, 0.75)$, and $F_X(x)$ is the CDF of $Beta(1, 2.5)$. **B**. 207 random uniform values from $Unif(0.35, 0.75)$. **C**. Beta distribution PDF with alpha = 1 and beta = 2.5.

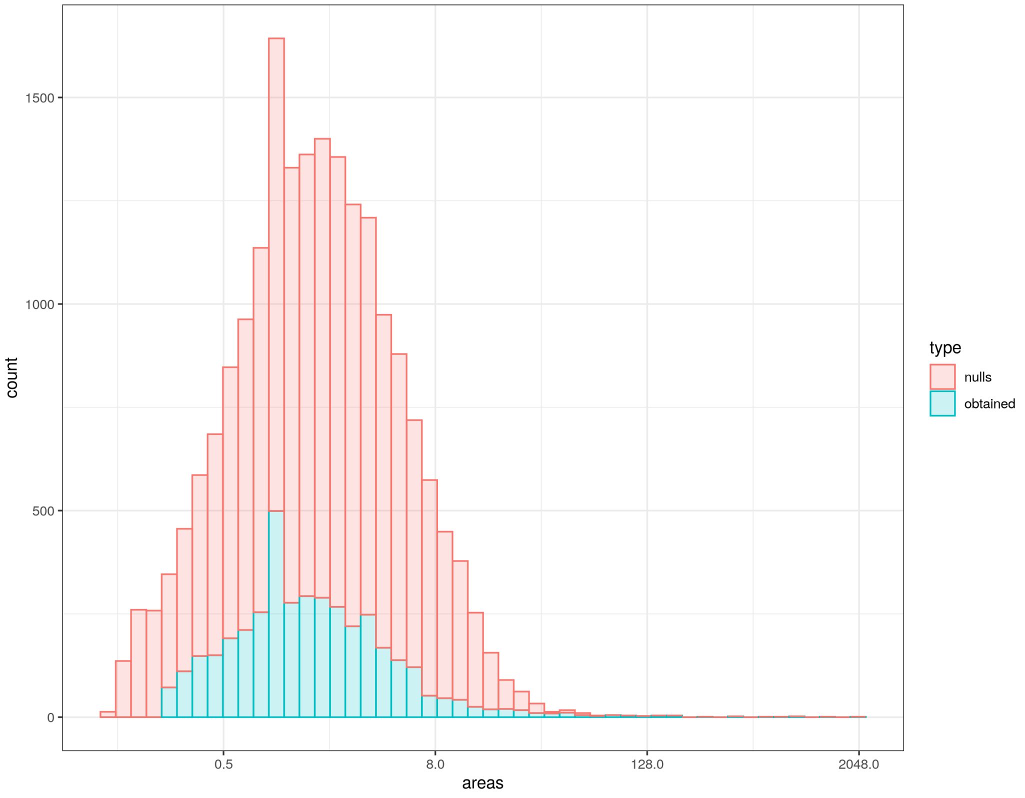


Supplementary Figure 10: Distribution of areas obtained from one of the 50 simulations by running the permutation method with $K = 0.2$. X-axis log2 transformed.
